# RNA-RNA Interactome Approaches Provide *in vivo* Evidence for a Critical Role of the Hfq Rim Face in sRNA-mRNA Pairing

**DOI:** 10.1101/2025.06.26.660752

**Authors:** Xing Luo, Aixia Zhang, Caroline Esnault, François De Mets, Philip P. Adams, Ryan K. Dale, Gisela Storz, Susan Gottesman

**Affiliations:** Laboratory of Molecular Biology, National Cancer Institute, Bethesda, MD 20892, USA; Division of Molecular and Cellular Biology, Eunice Kennedy Shriver National Institute of Child Health and Human Development, Bethesda, MD 20892, USA; Bioinformatics and Scientific Programming Core, Eunice Kennedy Shriver National Institute of Child Health and Human Development, Bethesda, MD 20892, USA; Biology of Spirochetes Unit, Laboratory of Bacteriology, Division of Intramural Research, National Institute of Allergy and Infectious Diseases, Bethesda, MD 20892, USA; NAV Drug Substance, GSK Vaccines, Cambridge, MA 02140, USA

## Abstract

Most bacterial small regulatory RNAs (sRNAs) modulate gene expression by forming complementary base pairs with target mRNAs, dependent upon the RNA chaperone Hfq. Hfq has three RNA-binding faces (proximal, rim, and distal), facilitating simultaneous binding of sRNAs and target mRNAs. Here, we systematically examined the functional impact of point mutations in the RNA binding faces on in vivo pairing, using the RNA Interaction by Ligation and Sequencing (RIL-seq) approach. The distal and proximal Hfq binding face mutants retained substantial numbers of RNA–RNA interactions (significant chimera counts or S-chimeras). However, the rim face mutant R16A showed a near-complete loss of S-chimeras, although Hfq R16A retained partial RNA binding activity as well as partial regulatory activity. Intracellular RIL-seq (iRIL-seq), a method with fewer *in vitro* processing steps, led to more S-chimeras in R16A but there were still fewer than for wild type Hfq and somewhat different sets of prevalent RNA-RNA pairs. Our analysis provides insights into how the RNA-binding faces of Hfq contribute to pairing *in vivo*, document the key role for the rim face in stabilizing RNA pairs on Hfq, and highlight intriguing differences captured by different RNA-RNA interactome approaches.

**GRAPHIC ABSTRACT:** 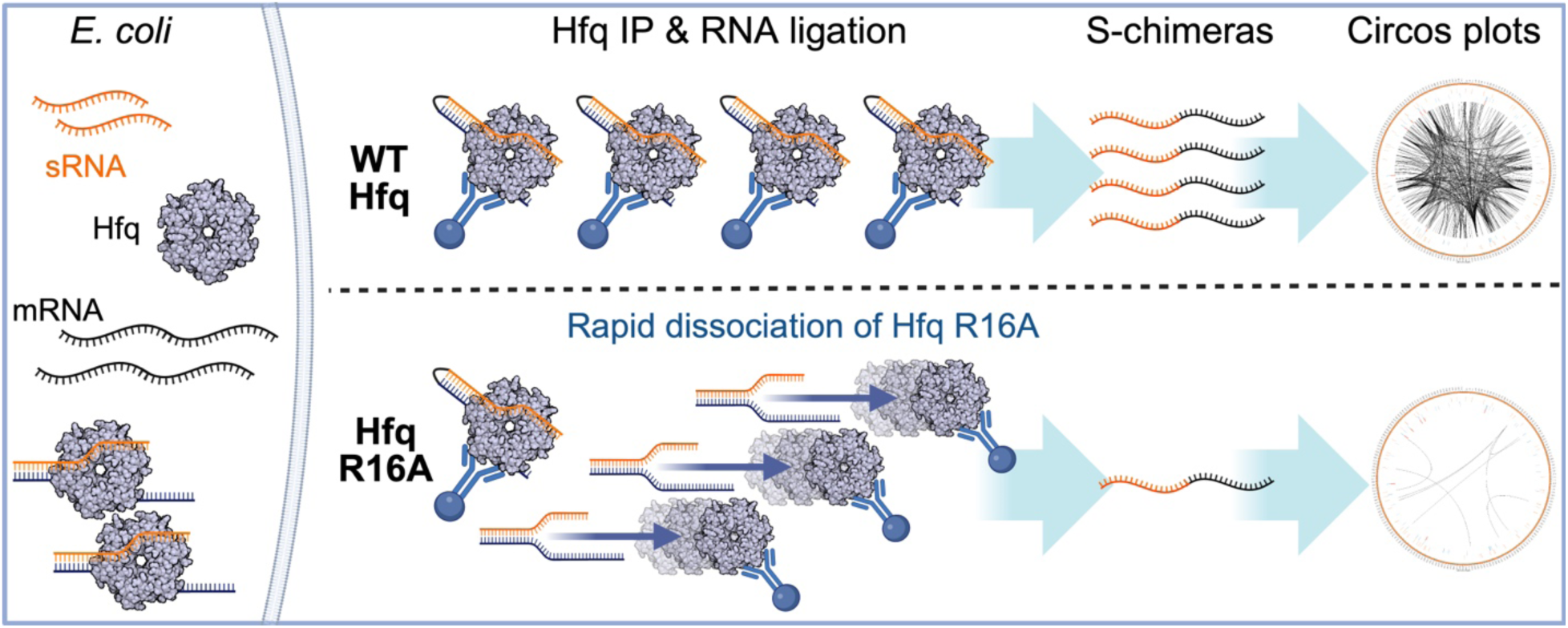

## INTRODUCTION

Gene expression in bacteria is finely tuned to adapt to changing environmental conditions. Central to this regulation are small RNAs (sRNAs), which modulate gene expression at the post-transcriptional level by base-pairing with specific mRNA targets (1–3). Depending on the target, sRNA binding can lead to translational inhibition, mRNA degradation, or in some cases, translational activation (1). This regulation plays critical roles in bacterial physiology, impacting stress responses, virulence, and metabolic pathways (4–6).

In *Escherichia coli* and many other bacterial species, the Sm-like RNA-binding protein Hfq acts as a chaperone to stabilize sRNAs and enhance their pairing with mRNA targets, making Hfq indispensable for many sRNA-mediated regulatory networks (7–9). Hfq forms a hexameric ring with three primary RNA-binding faces: the proximal face, distal face, and rim. Each face is specialized for binding specific RNA motifs, enabling Hfq to interact with multiple RNAs simultaneously (10,11). The proximal face generally binds U-rich sequences at the 3’ ends of sRNAs, aiding in their stabilization and facilitating interactions with mRNA targets (12,13). The distal face recognizes A-rich sequences in mRNA leaders, with a typical motif of ARN repeats, where A stands for adenine, R for any purine nucleotide, and N for any nucleotide (14–16). The rim, enriched with conserved arginine residues, plays a crucial role in binding and stabilizing certain sRNAs by interacting with UA-rich sequences (17,18). Recent *in vitro* studies have further shown that conserved arginine residues on the rim catalyze RNA base pair formation and exchange, supporting the RNA annealing activity of Hfq (19,20).

In Enterobacteriaceae, Hfq-bound sRNAs have been categorized into two main classes based on their interactions with the binding faces of Hfq; Class I sRNAs and Class II sRNAs (21). Class I sRNAs bind to the proximal and rim faces of Hfq, while their target mRNAs primarily bind to the distal face. In contrast, Class II sRNAs bind to the proximal and distal faces of Hfq, and their target mRNAs interact with the rim face (21).

Many studies on Hfq-RNA interactions have relied on *in vitro* approaches, including crystal structure analysis of Hfq/RNA complexes (13,14), stopped-flow FRET assays (17,19,20), size exclusion chromatography (13,18), and gel mobility shift assays using purified Hfq protein and RNA fragments (15,16). These studies have been instrumental in defining the RNA-binding and annealing properties of Hfq. However, these *in vitro* methods may not fully capture the operation of Hfq in a sea of cellular RNAs or the dynamic nature of Hfq’s function *in vivo*. *In vivo* characterization of Hfq mutants has been more limited. The functional consequences of Hfq face mutations were assessed using reporter assays or northern analysis to measure the effects of sRNA induction in different Hfq mutant backgrounds (12,21,22), as well as Hfq co-immunoprecipitation (IP) assays to examine Hfq-RNA interactions (22,23). These *in vivo* studies confirmed expected effects of Hfq face mutants on sRNA binding and stability. However, they primarily focused on a limited subset of model sRNAs and mRNAs, and many were carried out before we were aware of the two categories of Hfq-binding sRNAs.

In this study, we systematically investigated the contributions of individual Hfq faces to Hfq-RNA interactions and, more importantly, to sRNA-mRNA pairing, using RNA Ligation-Based RNA-seq (RIL-seq) approaches. RIL-seq is a powerful approach to capture *in vivo* RNA-RNA interactions mediated by RNA-binding proteins. The protocol involves UV crosslinking, protein IP, and *in vitro* RNase and RNA ligase treatments, which join two proximal RNA molecules into a chimeric RNA fragment. Both chimeric fragments and single RNAs are subsequently isolated and identified through deep sequencing (Fig. 1A) (24,25). To assess the contributions of the distinct RNA-binding faces of Hfq, we applied Total RNA-seq and RIL-seq to strains expressing chromosomal copies of Flag-tagged Hfq with single amino acid substitutions in the proximal, distal, or rim faces of Flag-tagged Hfq. Most mutants produced numbers of distinct RNA–RNA interactions with significant chimera counts (S-chimeras) comparable to those of wild-type Hfq. However, the R16A mutation on the rim face resulted in a near-complete loss of S-chimeras, pointing to a severe defect in Hfq’s ability to catalyze sRNA-mRNA pairing. We further investigated this observation to determine whether the lack of S-chimera in R16A reflects a limitation of the RIL-seq protocol or a specific defect in Hfq function *in vivo*. Our results reveal that the R16A mutation specifically impairs Hfq’s ability to bind some RNAs and more profoundly affects the stability of sRNA-mRNA pairing on Hfq, providing new insights into the mechanisms underlying the chaperone activity of Hfq. Additionally, our findings highlight how different interactome protocols capture different aspects of RNA-RNA base pairing interactions.

**Fig. 1.**
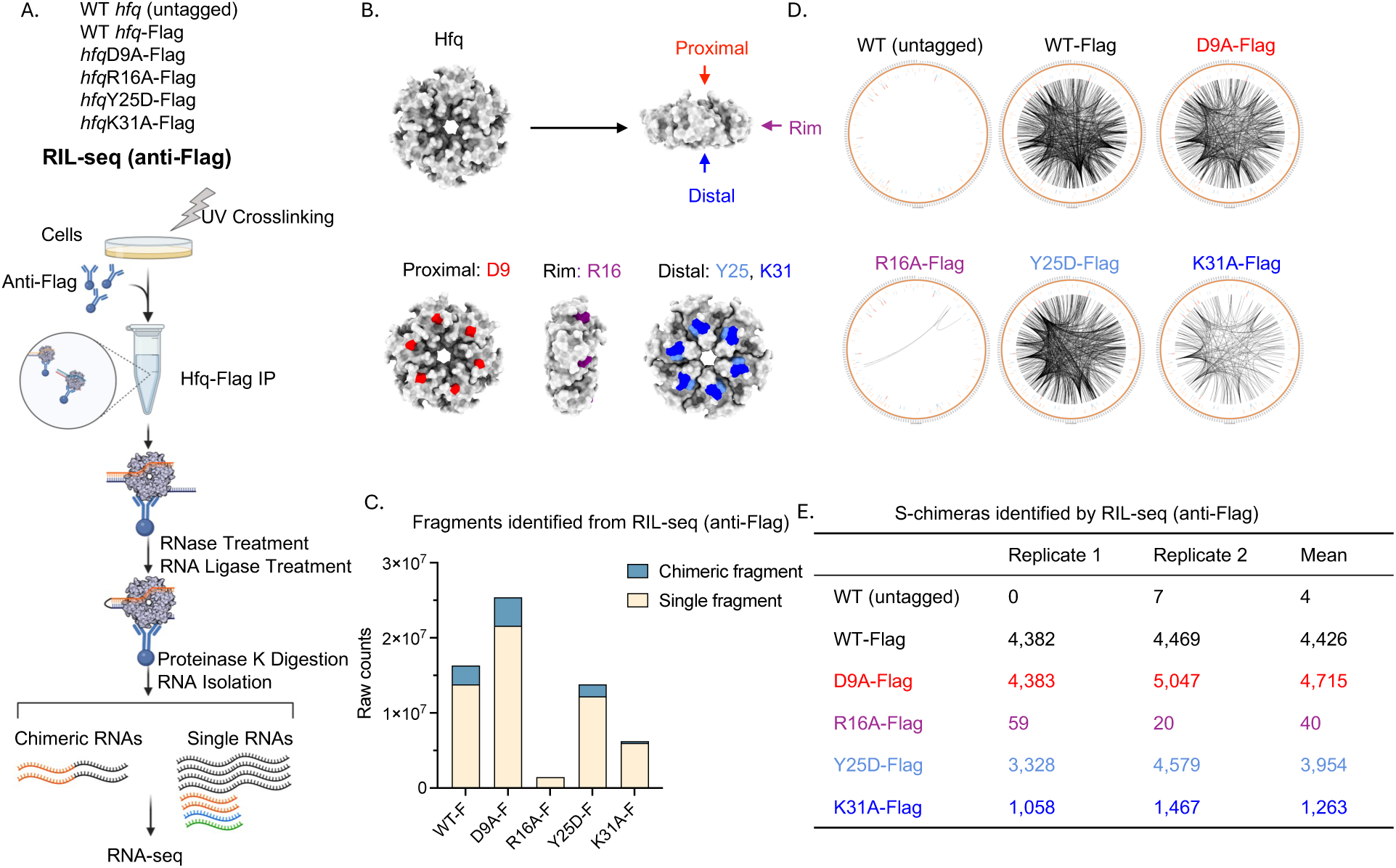
Hfq face mutants exhibit distinct effects on chimera formation. (A) Workflow of RIL-seq. Strains with 1X Flag tagged *hfq* (WT-Flag (FDM1004), D9A-Flag (FDM1005), R16A-Flag (FDM1006), Y25D-Flag (FDM1007), K31A-Flag (FDM1009)) and native *hfq* (Untagged MG1655)) were grown in LB (Lennox) medium at 37°C with shaking until an OD600 of 1.0 was reached. The cells were then harvested and processed for RIL-seq. Cell lysates were subjected to UV crosslinking, followed by Hfq immunoprecipitation (IP) using anti-Flag antibodies. After RNase and RNA ligase treatments, chimeric RNA fragments were isolated and sequenced, allowing identification of Hfq-dependent RNA-RNA interactions. sRNAs are shown in orange, mRNAs in black, and other Hfq-binding RNAs, such as tRNA and rRNA, are indicated in green and blue, respectively. This figure was created in BioRender (https://BioRender.com/jqqlsju). (B) Structural representation of the Hfq hexamer, highlighting the RNA binding faces and selected mutants. The Hfq hexamer has three RNA binding faces: the proximal face (red), rim face (purple), and distal face (blue). Mutants chosen to represent each binding face in this study include D9 from the proximal face, R16 from the rim face, and Y25 and K31 from the distal face. The structural models were generated using UCSF Chimera software (32). (C) Bar plot representing the number of single and chimeric RNA fragments identified from RIL-seq (anti-Flag). The counts represent the mean of two replicates, based on the data in Supplementary Fig. S3, with single fragments shown in beige and chimeric fragments in blue. (D) Circos plots displaying the RNA interactomes for each Hfq mutant as identified by RIL-seq. Each plot corresponds to one Hfq variant, with edges connecting genomic loci of interacting RNAs. The thickness of each edge reflects the normalized number of chimeric reads, indicating interaction strength. Circos plots were generated using Circos software (http://circos.ca/). (E) S-chimeras identified by RIL-seq (anti-Flag). Table displaying the number of RNA pairs with significant chimeric fragments (S-chimeras, chimeric fragments ≥ 5, *p* < 0.05, Fisher’s exact test) detected by RIL-seq (anti-Flag) for each Hfq mutant (Supplementary Table S4), with counts presented for two biological replicates and the calculated mean.

## MATERIALS AND METHODS

### Bacterial strains and plasmids

All bacterial strains used in this study were derived from *E. coli* K-12 strain MG1655 and are detailed in Supplementary Table S1, along with their sources or derivations. Mutant *hfq* alleles were first engineered in an intermediate strain (AZ233), which harbored a *miniλ*::*tet^R^*and *cat-sacB* cassette inserted at the *hfq* locus. This was achieved using λ-red recombineering with gBlocks synthesized by IDT (Supplementary Table S2). The modified *hfq* alleles were then introduced into the experimental strain NM572 via phage P1 transduction.

NM572 is a derivative of MG1655 provided by Dr. Nadim Majdalani and carries Δ*hfq*::*cat-sacB* and Δ*purA::kan* alleles. This strain served as the recipient for generating isogenic *hfq* mutant strains, both with and without a Flag tag (Supplementary Table S2). Donor strains carrying the desired *hfq* alleles were used for transduction, and recombinants were selected for *purA*^+^ restoration and screened for loss of the *cat^R^* gene. All genetic modifications were confirmed by DNA sequencing.

Plasmid pBAD24 (26) was from the lab collection. The T4 RNA Ligase I-expressing plasmid pBAD24-*t4rnl1* was synthesized by the GenScript company.

### RIL-seq (anti-Flag and anti-Hfq)

The strains were grown in LB (Lennox) at 37 °C with agitation. When the OD_600_ reached 1.0, 40 OD_600_ of cells were collected by centrifuging at 5000 rpm for 10 min at 4°C. RIL-seq was performed as previously described (24,25,27). The proteins and RNA molecules were crosslinked *in vivo* by exposing the cells to 800 mJ of 254 nm UV irradiation. After lysing the cells mechanically, Hfq and its bound RNAs were coimmunoprecipitated using either 3 µl of anti-Flag antibody (Sigma-Aldrich) or 100 µl of Hfq antibody (lab collection (28)). RNAs were trimmed by RNase A/T1 and then treated with T4 polynucleotide kinase (PNK) to generate fragments with 5’ P and 3’ OH ends that can be subsequently ligated. T4 RNA ligase was then used to ligate the neighboring RNAs. After digesting Hfq protein with proteinase K, the RNAs were isolated and subjected to sequencing library construction as described previously (27). The index primers are listed in Supplementary Table S2. Twelve cycles of PCR were performed at the last step to enrich each library. The libraries were sequenced by pair-end sequencing using Illumina NovaSeq. A step-by-step full protocol is available in the supplementary materials (Supplementary Protocol File 1).

### Total RNA-seq and RIP-seq

The strains were cultured in LB (Lennox) at 37 °C with agitation. When the OD_600_ reached 1.0, 20 OD_600_ of cells were collected by centrifuging at 5000 rpm for 10 min at 4°C. The cells were lysed by vortexing with 212-300 µm glass beads (Sigma) in a total volume of 2 ml lysis buffer (20 mM Tris-HCl/pH 8.0, 150 mM KCl, 1 mM MgCl_2_ and 1 mM DTT).

The isolation of total RNA and Hfq co-IP RNA was performed as previously described (29) with the following modifications: 1.9 ml of cell lysate were mixed with 100 µl of Hfq antibody and 120 mg of protein A-Sepharose CL-4B (GE Healthcare). Hfq co-IP RNA was isolated from protein A-Sepharose beads by extraction with phenol:chloroform:isoamyl alcohol (25:24:1), followed by ethanol precipitation. Total RNA was isolated from 100 µl of cell lysate by Trizol (ThermoFisher Scientific) extraction followed by chloroform extraction and isopropanol precipitation. RNA was dissolved in 15 µl of DEPC water.

cDNA libraries were constructed based on a protocol adapted to capture bacterial sRNAs too small to be captured by standard protocols, as well as mRNAs (29). 400 ng of total RNA samples or Hfq co-IP RNA samples were used as input and the index primers are listed in Supplementary Table S2. Seven cycles of PCR were performed at the last step to enrich each library. The sequencing was carried out using Illumina NovaSeq. The step-by-step full protocol is available in the supplementary materials (Supplementary Protocol File 2).

### Hi-GRIL-seq and iRIL-seq

Strains carrying either WT *hfq* (XL2001) or the *hfq* R16A mutant (FDM1050) were transformed with either an empty vector (pBAD24) or a T4 RNA Ligase I-expressing vector (pBAD24-*t4rnl1*). A single colony of each transformant was picked and inoculated in 10 ml of LB (Lennox) medium supplemented with 100 µg/ml ampicillin, then grown overnight at 37°C with agitation. The overnight cultures were diluted into 80 ml of fresh medium to an OD_600_ of 0.05 and incubated at 37°C with agitation. When the OD_600_ reached 0.5, 0.2% arabinose was added to induce T4 RNA Ligase I expression. Cultures were further grown until the OD_600_ reached approximately 1.0, at which point 20 OD_600_ units of cells were collected by centrifugation at 5000 rpm for 10 minutes at 4°C. The cells were lysed by vortexing with 212-300 µm glass beads (Sigma) in a total volume of 2 ml lysis buffer (20 mM Tris-HCl/pH 8.0, 150 mM KCl, 1 mM MgCl_2_ and 1 mM DTT).

Isolation of total RNA and Hfq co-IP RNA was performed as described in the Total RNA-seq and RIP-seq sections above. For each library, 400 ng of total RNA or Hfq co-IP RNA was used as input for cDNA synthesis, with total RNA serving as input for Hi-GRIL-seq and Hfq co-IP RNA for iRIL-seq. Library construction followed the same procedure outlined in the Total RNA-seq and RIP-seq sections, except that ribosomal RNA was depleted using DASH, adapted from (30) and described in detail below, rather than Ribo-Zero. Index primers used for library preparation are listed in Supplementary Table S2. Final enrichment of libraries was performed with seven cycles of PCR. Sequencing was conducted on the Illumina NovaSeq platform. The step-by-step full protocol is available in the supplementary materials (Supplementary Protocol File 3).

### rRNA depletion using the DASH protocol

For cDNA library construction in RIL-seq, RIP-seq, and Total RNA-seq experiments, ribosomal RNA was removed using the Ribo-Zero rRNA Removal Kit for bacteria (Illumina, #MRZGN126). After the discontinuation of this kit, we adapted DASH (depletion of abundant sequences by hybridization) (30) to remove *E. coli* rRNA for Hi-GRIL-seq and iRIL-seq. DASH uses the Cas9 nuclease and guide RNAs (gRNAs) to target Cas9-mediated cleavage prior to library amplification.

Target sequences within *E. coli* rRNA genes were identified using a custom algorithm based on https://github.com/NICHD-BSPC/bacteria-dash. Briefly, rRNA regions coordinates were identified from the annotation EcoCyc version 19.0 (http://ecocyc.org) (31) and represented the regions to deplete. The *E. coli* reference genome (GCF_000005845.2, ASM584v2) was scanned for all Cas9 PAM sites (NGG) on both strands adjacent to the regions to deplete. The candidate sgRNAs were defined as the 20 bp regions adjacent to the 5’ end of PAM sites. These candidates were filtered based on the following thresholds: a GC content > 30% and < 80%, Tm threshold of heterodimer formation with the primer of 40, and end stability Tm threshold of 30. Off-target candidates were defined as mapping to regions outside of the target regions in addition to the target regions and were discarded. This resulted in a list of 822 sequences (Supplementary Table S2), which were purchased from IDT as a unique oligo pool, each with the following structure (5′ to 3′): T7 promoter (TTCTAATACGACTCACTATA) + gRNA sequence + scaffold (GTTTTAGAGCTAGAAATAGC). Single guide (sg) RNA pools were *in vitro* transcribed as previously described (30).

RNA-seq library construction started by following the initial steps of the bacterial-sRNA adapted, RNAtag-seq methodology (27). RNA samples were fragmented, DNase and FastAP (Thermosensitive Alkaline Phosphatase) treated, and subjected to RNA 3′ adaptor ligation. Immediately following the 3′ adaptor ligation, samples were pooled together and cleaned using the RNA Clean and Concentrator-5 kit (Zymo) according to the manufacturer’s instructions to a final volume of 13 μl in RNase-free water. Next, 11 μl of the eluted sample was subjected to first-strand cDNA synthesis, RNA degradation, RNACleanXP bead clean-up, cDNA 3′ adaptor ligation, and two rounds of RNACleanXP bead clean-up, following the steps of the RNAtag-seq methodology but skipping Ribo-Zero steps (27). The cDNA was eluted from the RNACleanXP beads with 18 μl RNase-free water. Eight microliters of cDNA were pre-amplified with 2 cycles of PCR using KAPA HIFI and the P5_Enr primer and P7_BC1_Enr primer. Pre-amplified PCR products were cleaned with RNACleanXP beads and quantified by NanoDrop analysis (the typical concentration is 10-20 ng/μl). Cas9 was pre-loaded with *E. coli* sgRNAs, incubated at 37°C for 2 h with the pre-amplified cDNA at a ratio of 1:3:6 (cDNA:Cas9:sgRNA), and subsequently treated with Proteinase K and PMSF, as previously described (30). The amounts of cDNA, Cas9, and sgRNA can be calculated using the DASH calculator, provided in the supplementary materials (Supplementary Protocol File 4). rRNA-cut cDNA was cleaned with RNACleanXP beads and PCR amplified prior to RNA-sequencing, following the PCR enrichment test and QC analysis for RNAtag-seq (27). The resulting iRIL-seq and Hi-GRIL-seq libraries prepared with the modified DASH protocol resulted in ∼ 10% sequencing reads to rRNA genes, similar to results obtained using the Ribo-Zero rRNA Removal Kit for bacteria used for RIL-seq, RIP-seq and Total RNA-seq experiments.

### Construction of a comprehensive *E. coli* genome annotation file

Analysis of genome-wide RNA expression or other approaches is frequently done by assigning signals to annotated features (genes), but this can lead to missing interesting and biologically relevant signals from previously unrecognized transcribed regions of the chromosome. A comprehensive *E. coli* genome annotation file to simplify the identification of RNA features was constructed using Snakemake v7.25.0. The foundation for this file was the annotation from previous papers (24,25,27), which included mRNAs, sRNAs, and known UTRs from EcoCyc version 19.0 (http://ecocyc.org) (31). Predicted UTRs were defined by extending 100 nucleotides upstream of each start codon (ATG) and downstream of each stop codon, unless these extensions overlapped another transcript, in which case shorter lengths were used. Antisense (AS) regions were designated as sequences located on the strand opposite to annotated coding regions. Intergenic regions (IGRs) were identified in a strand-specific manner as the sequences separating any annotated features on the same strand. Transcription unit intergenic regions (TUs) were defined as inter-feature regions less than 100 base pairs in length (Supplementary Fig. S1). This base annotation was updated by integrating the NC_000913.3_gene_tRNA.gff file (28) and incorporating known sRNA coordinates reported in a recent review (1). These datasets were merged as pandas DataFrames, and antisense and intergenic regions were recalculated using pybedtools v0.9.0. To enrich the annotation with gene functions, gene descriptions were added from a SmartTable export from EcoCyc (accessed on 06/22/2023). Feature names were standardized by combining the feature type (“UTR”, “AS”, “IGR”, or “TU”) with adjacent gene names, separated by periods; for IGRs and TUs flanked by two genes, both gene names were included (e.g., gene1.gene2.featuretype). The final annotation file, named NC_000913-3_all_feature.gff, was used for all RNA-seq analysis in this study and is available in GEO alongside the raw sequencing data.

### RIL-seq data processing

RNA–RNA interactions were detected by identifying chimeric reads within RIL-seq libraries. Raw FASTQ files were processed through a Snakemake-based pipeline incorporating tools from lcdb-wf v1.10.2 (https://github.com/lcdb/lcdb-wf) and RIL-seq v0.82 (https://github.com/asafpr/RIL-seq). Initial demultiplexing was performed using cutadapt v4.3 with parameters “--error-rate=0.2 --overlap=9”. Adapter trimming and quality filtering were subsequently applied using cutadapt with the parameters “--nextseq-trim 20 --overlap 6 --minimum-length 25”. Trimmed reads were aligned to the *E. coli* reference genome (GCF_000005845.2, ASM584v2) using the RIL-seq script “map_single_fragments.py”, with the parameters “--feature exon --create_wig”.

Chimeric reads were identified using “map_chimeric_fragments.py” with the “-r” parameter to account for the reverse-stranded library design. Significant RNA–RNA interaction regions were then identified using “RILseq_significant_regions.py”, with the parameters “--total_reverse”, “—ribozero”, and “--BC_chrlist “chr”” to specify strand orientation, exclude rRNA reads, and restrict analysis to the main chromosome, respectively. Chimeras outside of statistically significant regions were filtered as non-specific and excluded from further analysis.

Significant chimeric fragments were assigned to overlapping genes in R v4.2.2 using GRanges objects from the GenomicRanges v1.50.0 package. Overlap was determined using the findOverlaps function from IRanges v2.32.0. When fragments overlapped multiple genes, a fragment-to-gene ratio was calculated; if a single gene accounted for more than 70% of overlapping reads (ratio > 0.7), it was assigned exclusively, otherwise all overlapping genes were retained.

### Total RNA-seq and RIP-seq processing

Total RNA-seq and RIP-seq samples were processed using the lcdb-wf v1.10.2 pipeline (https://github.com/lcdb/lcdb-wf). Raw FASTQ files underwent demultiplexing, adapter trimming, and quality filtering with cutadapt v4.3, first using the parameters “--error-rate=0.2 --overlap=9”, followed by “--nextseq-trim 20 --overlap 6 --minimum-length 25”. Trimmed FASTQ files were mapped to the *E. coli* reference genome (GCF_000005845.2, ASM584v2) using BWA v0.7.17 with default parameters.

Gene expression was quantified using featureCounts from Subread v2.0.3 with the “-0” parameter enabled to allow counting of reads in overlapping features, using the NC_000913-3_all_feature.gff annotation file. Differential expression analysis was performed in R v4.2.2 using DESeq2 v1.38.0. For RIP-seq samples, enrichment ratios were computed by contrasting IP samples with their corresponding total RNA levels.

### Northern blot

The northern blot was performed based on the method described in (29). Briefly, 10 µg of total RNA or 1 µl of Hfq co-IP RNA were resolved in 8% polyacrylamide urea (6M) gels and then transferred to Zeta-Probe GT membranes (Bio-Rad) and crosslinked by UV irradiation. After being blocked in ULTRAhyb-Oligo Hybridization buffer (Ambion) for 2 h at 45 °C, the membranes were incubated overnight with probes (listed in Table S4) which have been prelabeled by ^32^P at the 5’ end. The membranes were washed twice with 2× SSC/0.1% SDS and once with 0.2× SSC/0.1% SDS at room temperature, 25 min with 0.2× SSC/0.1% SDS at 45 °C, followed by a final wash with 0.2× SSC/0.1% SDS at room temperature. The probing signals were measured by exposing to X-ray films.

### Western blot

In general, 1 ml of bacterial culture was centrifuged at 4 °C for 2 minutes, and the resulting pellet was rapidly frozen in liquid nitrogen. For every 1 ml of cells at OD_600_ ∼ 1.0, 100 µl of 1× LDS (lithium dodecyl sulfate) sample buffer (ThermoFisher Scientific, #84788) and 50 mM dithiothreitol (DTT) were added to dissolve the protein. A total of 10 µl boiled samples were resolved on a 4 -12% Bis-Tris NuPAGE protein gel (Invitrogen, NP0323BOX) in 1× MES (2-(N-morpholino)ethanesulfonic acid) buffer (Invitrogen) for 50 min at 150 V. Resolved proteins were transferred to a nitrocellulose membrane using iBlot 2 Dry Blotting System (ThermoFisher Scientific). The membranes were blocked with 5% nonfat milk in PBST buffer (1X PBS and 0.03% Tween-20) 1 hour at room temperature (RT), then incubated with primary antibodies and fluorescent secondary antibodies at RT for 1 h each. Primary antibodies used in this study were lab collection polyclonal rabbit anti-Hfq antibody (1:5000), polyclonal rabbit anti-RpoS antibody (1:5000), monoclonal anti-Flag M2-alkaline phosphatase antibody (1:3000; Sigma, #A9469), mouse monoclonal anti-6X His tag antibody (1:2000, abcam, #ab117504) and monoclonal mouse anti-EF-Tu antibody (1:10000; LifeSpan BioSciences Inc.). Fluorescent secondary antibodies were StarBright Blue 700 goat anti-rabbit (Bio-Rad) and/or Dylight 800 goat anti-mouse (Bio-Rad). Before and after secondary antibodies, membranes were washed 3× 10 min with PBST buffer. Fluorescent signals were measured by the ChemiDoc MP imaging system (Bio-Rad) and quantified with ImageJ Software.

## RESULTS

### Hfq face mutants differentially impact Hfq-associated chimera formation

To explore the roles of each RNA-binding face of Hfq in RNA binding and pairing, we selected four representative single amino acid substitutions in the proximal, rim, and distal faces. These mutations were chosen based on insights from previous studies, particularly the work in (22), which tested 14 Hfq mutants for their ability to mediate regulation by seven sRNAs. The proximal face mutant D9A (Fig. 1B) was included based on observations of both positive and negative roles of this residue in *in vivo* regulation, as well as *in vitro* studies (17). The rim face mutant R16A (Fig. 1B) was chosen because mutations of R16, a highly conserved residue, have been shown to partially impair Hfq function (19). Similarly, the distal face mutant Y25D (Fig. 1B) was included because it also partially disrupts Hfq function, with some sRNAs losing activity with others remaining unaffected. In contrast, K31A, another distal face mutant (Fig. 1B), was selected due to its relatively mild impact on Hfq function, allowing us to compare the effects of mutations with differing levels of disruption (22).

In the original RIL-seq protocol (27), a Flag tag is fused to the RNA-binding protein to facilitate efficient IP. To this end, we generated chromosomal *hfq* mutants with a single C-terminal Flag tag and included a native untagged strain as a negative control for the IP. Western blots using an anti-Hfq antibody indicated that the Flag tag slightly reduced Hfq levels (Supplementary Fig. S2A, compare lane 3 to lane 1). The R16A-Flag mutant was poorly detected by the anti-Hfq antibody but was readily detected by the anti-Flag antibody (lane 5), suggesting that the combination of the Flag tag and R16A mutation interfered with epitope recognition by the anti-Hfq antibody. We performed RIL-seq with Flag-tagged wild-type Hfq (WT-Flag), the *hfq* point mutants (D9A-Flag, R16A-Flag, Y25D-Flag, and K31A-Flag), and an untagged wild-type Hfq (WT) strain. In parallel, we also performed Total RNA-seq to evaluate the gene expression profiles in the mutant backgrounds. Biological duplicates of all strains were grown in LB (Lennox) medium at 37°C to early stationary phase (OD_600_ ∼ 1.0). When analyzing all the deep sequencing data, we used a newly created annotation file (NC_000913-3_all_feature.gff), derived from a previously published annotation (28), to ensure comprehensive *E. coli* genome coverage and to better define signals for all RNA fragments. This updated annotation extends beyond known open reading frames (ORFs) and non-coding RNAs to incorporate intergenic regions (IGRs), estimated untranslated regions (EST5UTR and EST3UTR) and antisense RNAs (AS), resulting in a total of 20,320 annotations across the genome (Supplementary Fig. S1 and Materials and Methods). Thus, the annotation file captures a broader range of genomic features, including previously unannotated regions, allowing for a more complete analysis of RNA interactions.

Principal Components Analysis (PCA) of the Total RNA-seq results (Supplementary Table S3) revealed high reproducibility between replicates (Supplementary Fig. S2B). WT and WT-Flag samples clustered closely together (equating to lower variation among the datasets; Supplementary Fig. S2B), with only 2.5% of genes differentially expressed due to the addition of the Flag tag; this set did not include many sRNAs (Supplementary Fig. S2C, Table S3). Therefore, we conclude that the Flag tag does not have a major effect on Hfq function. The overall effects of Hfq point mutations on global gene expression varied. The K31A-Flag samples clustered closest to WT-Flag (Supplementary Fig. S2B), with only 4.8% of genes differentially expressed compared to WT-Flag (Supplementary Fig. S2C and Supplementary Table S3). This aligns with previous findings that K31 mutations minimally disrupt Hfq activity (22). The other mutations led to more genes differentially expressed, with greater separation from WT-Flag in the PCA plot (Supplementary Fig. S2B). Differential gene expression analysis of R16A-Flag, D9A-Flag, and Y25D-Flag showed 7.7%, 13.9% and 17.2% of genes, respectively, differentially expressed compared to WT-Flag (Supplementary Fig. S2C and Supplementary Table S3).

In RIL-seq, the chimeric RNAs reflect RNA-RNA interactions facilitated by Hfq. With WT-Flag, approximately 15% of the total identified RNA fragments were chimeras, while the remainder were single RNA fragments (Fig. 1C, Supplementary Fig. S3). The D9A and Y25D mutants had similar levels of chimeric RNA, with 15% and 11%, respectively. In contrast, for K31A-Flag and R16A-Flag, we identified substantially fewer total RNA fragments (single + chimeric RNA fragments), and among these fragments, the percentage of chimeras was also lower, approximately 4% and 1%, respectively (Fig. 1C, Supplementary Fig. S3). Using a previously published algorithm (24), we identified RNA-RNA interactions supported by significant chimeras (S-chimeras), where each S-chimera represents a different pair of RNAs. These were considered significant based on a threshold of ≥ 5 chimeric fragments and p < 0.05 (Fisher’s exact test) for each strain (Supplementary Table S4). These interactions were visualized using Circos plots (http://circos.ca/) (Fig. 1D), where lines connect the genomic loci of interacting RNAs, representing RNA–RNA interactions. The numbers of S-chimeras are summarized in Figure 1E, showing distinct effects of some of the Hfq mutants on the numbers of RNA-RNA interactions. As expected, almost no detectable S-chimeras were identified in the untagged WT strain, consistent with the S-chimeras resulting from Hfq-facilitated interactions. In the WT-Flag strain, around 4,400 S-chimeras were identified (Fig. 1E). The D9A-Flag mutant produced a comparable number of S-chimeras to WT Hfq, suggesting minimal impact of the D9 proximal face mutation on S-chimera formation ability. The role of D9 in Hfq function is not well understood (17). While D9A formed a similar level of S-chimeras as WT, the pattern for both total RNA and S-chimeras differed from WT and requires more experimentation and analysis to interpret. Here, we focused on the other Hfq mutants. The distal face mutants, Y25D-Flag and K31A-Flag, exhibited reduced S-chimera counts (average numbers of 3,954 and 1,263, respectively), with K31A having the most pronounced reduction, despite previous observations that K31A causes minimal effects on Hfq-dependent regulation (22) (Fig. 1D, 1E; total RNA levels, Supplementary Fig. S2B, C). Most strikingly, the R16A-Flag mutant, which disrupts the rim face of Hfq, exhibited a near-complete loss of S-chimeras (average of 40, a 100-fold decrease compared to WT-Flag), indicating a severe defect in chimera formation. Capture of single fragments was also decreased for R16A, but only by 10-fold (Fig. 1C; Supplementary Fig. S3).

To assess whether the few S-chimeras associated with R16A-Flag corresponded to the most abundant interactions detected with WT-Flag, we chose the top 25 most abundant WT-Flag S-chimeras and determined their presence and abundance in R16A-Flag, Y25D-Flag, and K31A-Flag. Only six of the top 25 S-chimeras were detected in both R16A mutant datasets (Supplementary Table S5, Top). In contrast, most of the top WT-Flag S-chimeras were detected for Y25D-Flag and K31A-Flag, though with some differences in rank. In a reciprocal analysis for the top 20 S-chimeras for R16A-Flag, several were not detected in either WT-Flag dataset, suggesting that these S-chimeras reflect abnormal binding or interactions not normally promoted by Hfq (Supplementary Table S5, Bottom, marked as ND in column K or M).

### Hfq face mutants differentially impact chimera formation of both Class I and Class II sRNAs

To further investigate the effects of the mutations on specific sRNAs, we selected six representative, previously-studied sRNA-mRNA pairs for detailed analysis. We analyzed three Class I sRNA-mRNA pairs (Fig. 2A, top): MicA-*ompA* (33,34), DsrA-*rpoS* (35,36), and GcvB-*dppA* (37), and three Class II sRNA-mRNA pairs (Fig. 2A, bottom): ChiX-*chiP* (38), MgrR-*eptB* (39,40), and CyaR-*ompX* (41). The selected sRNAs were highly expressed under the experimental conditions (Supplementary Table S3). The target mRNAs for these sRNAs were selected based on three criteria: (1) high expression levels under the conditions of our experiments, as indicated by Total RNA-seq data (Supplementary Table S3), (2) chimera numbers among the highest for their respective sRNAs, as shown in RIL-seq data (Supplementary Table S4, Hfq-FLAG tabs), and (3) previous characterization in published studies. Most of these sRNAs function as negative regulators of their target mRNAs. The exception is DsrA, which positively regulates RpoS translation by base-pairing with the long mRNA 5’ UTR. This interaction disrupts a repressive secondary structure, thereby exposing the ribosome binding site and facilitating translation initiation (35,36). In previous work, we found that Class I sRNAs bind to the proximal face and rim of Hfq, and become more unstable in rim mutants; they are stabilized by mutations in the distal face that block binding of their target mRNAs; Class II sRNAs bind to the proximal and distal faces, and are thus destabilized by distal face mutants, while they are stabilized by rim mutations (21).

**Fig. 2.**
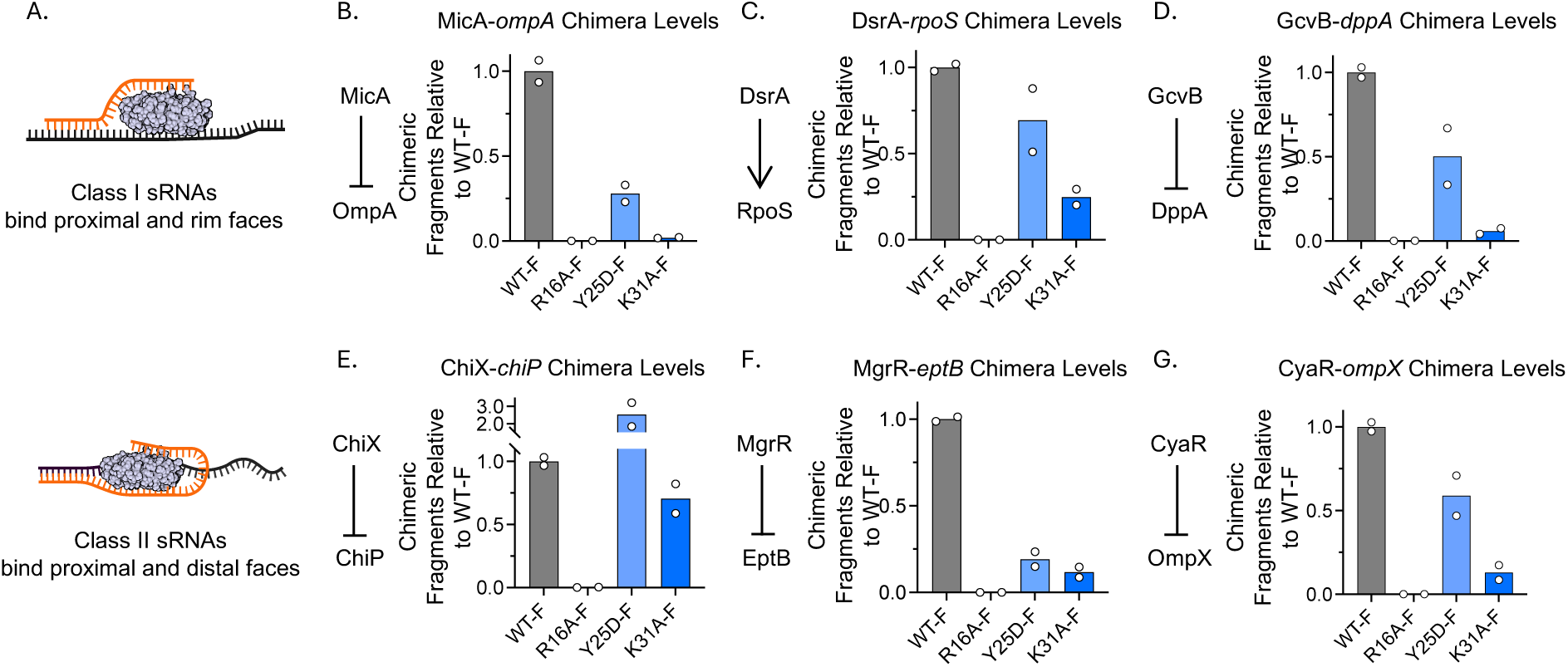
Hfq face mutants exhibit distinct effects on chimera formation of both class I and class II sRNAs. (A) Schematic representation of Class I and II sRNAs (orange) binding to the proximal and rim faces of Hfq, with target mRNAs shown in black. This figure was created in BioRender (https://BioRender.com/jqqlsju). (B–G) Quantification of chimera levels for Class I and Class II sRNA-mRNA pairs in *hfq* mutants, as measured by RIL-seq (anti-Flag). Note that R16A is a mutant on the rim, Y25D and K31A are on the distal face. Panels show representative interactions for Class I sRNAs: (B) MicA-*ompA* (33,34), (C) DsrA-*rpoS* (35,36) and (D) GcvB-*dppA* (37); and Class II sRNAs: (F) ChiX-*chiP* (38), (G) MgrR-*eptB* (40) and (H) CyaR-*ompX* (41). A full dataset of RIL-Seq results is available in Supplementary Table S4, where each tab corresponds to a replicate from a specific strain. To retrieve data for specific sRNA–mRNA interactions, refer to the RNA1_name (Column F) and RNA2_name (Column G) columns. Typically, the sRNA is listed in RNA2, and its target mRNA 5′UTR appears in RNA1 (e.g., ompA.5UTR, dppA.5UTR). Note: For *rpoS*, whose long 5′UTR overlaps with the upstream gene *nlpD* (42), the DsrA target is annotated as *nlpD* in the RNA1_name field. The number of identified chimeric fragments for each sRNA–mRNA pair is provided in Column P (interactions). Chimera counts for each mutant were normalized to the WT-Flag strain and are displayed as fold-change ratios. Bars represent the average of two biological replicates, with individual replicate values shown as white dots.

First, we examined the levels of the six representative sRNAs and their corresponding target mRNAs from Total RNA-seq data, normalizing them to WT-Flag levels (Supplementary Fig. S4 and Supplementary Table S3). Overall, the effects of the mutants on these sRNAs were consistent with previous work (21). The levels of Class I sRNAs decreased in the rim face R16A-Flag mutant and increased in the Y25D-Flag and K31A-Flag distal mutants (Supplementary Fig. S4A-C, left panels). Class II sRNAs remained largely unchanged in the R16A-Flag mutant and had reduced levels in the distal face mutants (Supplementary Fig. S4D-F, left panels). The levels of most of the target mRNAs were unaffected by mutations in Hfq, remaining either unchanged or somewhat elevated compared to WT-Flag (Supplementary Fig. S4A-F, right panels). The most dramatic changes were increases seen for two targets that are under strong Class II sRNA negative regulation, *chiP* (38) and *ompX* (41).

We then analyzed the interactions of the six sRNA-mRNA pairs (Fig. 2) using the RIL-seq data (Supplementary Table S4). The chimeric RNA levels of each sRNA-mRNA pair, extracted from Supplementary Table S4 (Column P), were normalized to WT-Flag, generating fold-change ratios shown in Fig. 2. In general, both Class I and Class II sRNAs showed similar patterns of relative chimera numbers compared to WT-Flag across the different Hfq mutants. The relative numbers of chimeras for both sRNA classes were drastically decreased in the R16A-Flag mutant compared to WT-Flag (Fig. 2B–G), aligning with the near-complete loss of total chimera amounts shown in Figure 1D and 1E, even though the total RNA levels of Class II sRNAs in the mutant were not notably reduced (Supplementary Fig. S4D-F, left panels). The distal face mutants Y25D-Flag and K31A-Flag had reduced chimera numbers for most sRNA-mRNA interactions (Fig. 2B–G), with K31A-Flag consistently displaying the most pronounced reduction. This result aligns with the total chimera counts observed in the distal face mutants, with a modest decrease for Y25D-Flag (90% of WT-Flag) and a more pronounced decrease for K31A-Flag (28% of WT-Flag) (Fig. 1D and 1E).

### Regulatory activity of Hfq mutants does not fully align with RIL-seq chimera results

We compared the S-chimera numbers identified for each Hfq mutant (Fig. 1E) to findings from our previous study on Hfq activity in binding face mutants (22). In the previous study, the K31A mutation was reported to have a mild impact on Hfq function, measured with reporter fusions, while the Y25D mutation was more defective. However, in the RIL-seq data, K31A-Flag exhibited a significant reduction in S-chimera numbers compared to Y25D-Flag (Fig. 1E). Additionally, the R16A-Flag mutation, which resulted in an almost complete loss of S-chimeras in our RIL-seq analysis (Fig. 1E), was still functional for some assays of regulatory activity in the prior study (22). The earlier study (22) utilized multi-copy plasmids to express sRNAs, whereas the RIL-seq experiments were performed with sRNAs expressed from the chromosome without overexpression. It is possible the higher levels of sRNAs partially suppressed the defects in these Hfq mutants, increasing the likelihood of productive interactions on Hfq. However, even with sRNAs expressed from the chromosome, our total RNA data suggested that Y25D-Flag is more defective than K31A-Flag and R16A-Flag, (Supplementary Fig. S2B, C), even though K31-FLAG and R16A-FLAG have fewer S-chimeras.

To investigate whether the significant changes in S-chimera number identified from RIL-seq reflect differences in the ability of sRNAs expressed at native levels to regulate their targets, we examined the expression of RpoS, which is positively regulated by sRNAs DsrA, RprA, and ArcZ under various conditions (Fig. 3, left panel) (43). The genome browser view generated from RIL-seq data showed that these sRNAs base-paired with the long 5′ UTR of *rpoS*, which overlaps with the upstream gene *nlpD* (42) (Fig. 3A, right panel). In WT-Flag, this region displayed a strong interaction signal, which was greatly reduced in K31A-Flag and nearly absent in R16A-Flag, while Y25D-Flag showed only a modest decrease (Fig. 3A, right panel). Chimera numbers of the three sRNA– *rpoS* pairs reflected this trend, with WT-Flag showing the highest interaction levels, R16A-Flag the lowest, and Y25D-Flag retaining more interactions than K31A-Flag (Fig. 3B).

**Fig. 3.**
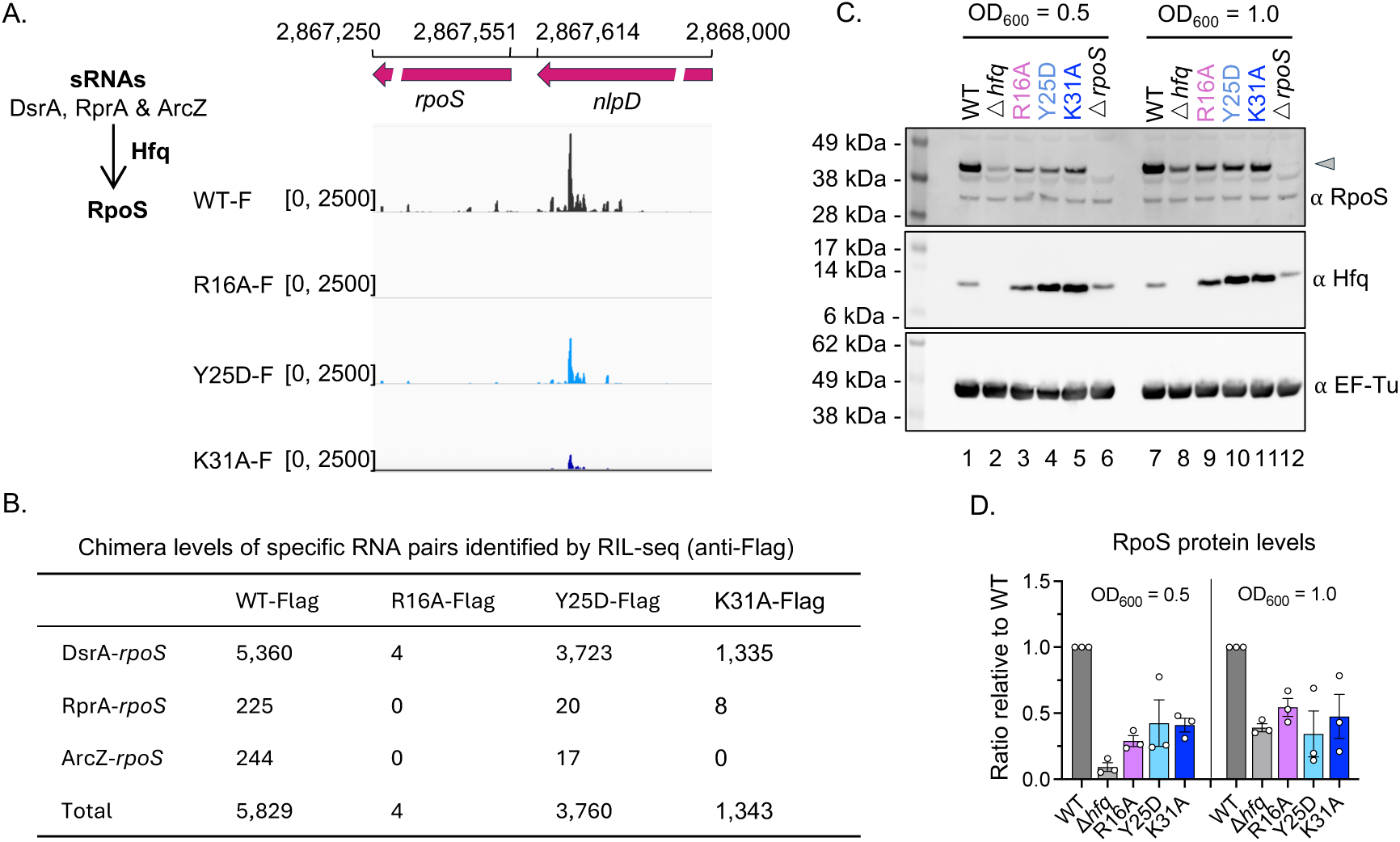
RIL-seq chimera counts do not fully reflect Hfq’s role in sRNAs regulation of RpoS levels. (A) Genome browser view (right panel) of the *rpoS* 5′UTR region targeted by sRNAs (left panel), as identified by RIL-seq (anti-Flag) in WT-Flag, R16A-Flag, Y25D-Flag, and K31A-Flag strains. The displayed region spans coordinate 2,867,250 to 2,868,000 on the *E. coli* chromosome. The coordinates marking the end of the *nlpD* coding region (2,867,614) and the start of the *rpoS* coding region (2,867,551) are indicated. (B) Chimera levels of DsrA-*rpoS*, RprA-*rpoS* and ArcZ-*rpoS* identified by RIL-seq (anti-Flag) in each strain. Chimera levels were extracted from Supplementary Table S4, following the same approach as described for Figure 2B, and are presented as the mean of two replicates. Note: *rpoS* 5′UTR overlaps with the upstream gene *nlpD* (42), and is therefore annotated as “nlpD” in the RNA1_name field. (C) Western blots for RpoS, Hfq, and EF-Tu (loading control) in WT (XL2001), Δ*hfq* (XL2002), *hfq* R16A (FDM1050), *hfq* Y25D (XL2003), *hfq* K31A (XL2003) and Δ*rpoS* (XL2019). Strains were grown in LB (Lennox) medium at 37°C with shaking, and 1 ml of cells was collected at OD600 values of 0.5 and 1.0 for western blot analysis. Molecular weight markers (kDa) are indicated on the left. (D) Quantification of RpoS Protein Levels from three independent experiments of 3C. Western blot bands were quantified using ImageJ software.

We measured RpoS protein levels by Western blot in six strains—WT *hfq*, τΔ*hfq*, *hfq* R16A, *hfq* Y25D, *hfq* K31A, and a *rpoS* deletion strain—grown to OD_600_ values of 0.5 and 1.0 (Fig. 3C). In the WT *hfq* strain, RpoS was strongly expressed under both conditions due to positive regulation by sRNAs (Fig. 3C, lanes 1 and 7). In the τΔ*hfq* strain, where sRNA-mediated positive regulation is impaired, RpoS was barely detectable at OD_600_ 0.5 (Fig. 3C, lane 2; Fig. 3D) and only weakly expressed at OD_600_ 1.0 (Fig. 3C, lane 8; Fig. 3D). Interestingly, all three *hfq* mutants showed similar intermediate RpoS levels at both ODs—higher than in Δ*hfq* but lower than in WT— despite showing markedly different chimera counts (Fig. 3A-B). This suggests that R16A retains partial function in mediating sRNA regulation of RpoS, which is not reflected in its near-complete loss of S-chimeras in RIL-seq (Fig. 1E). Additionally, although Y25D-Flag exhibited substantially more chimeras (Fig. 3A-B), its RpoS levels were comparable to those of R16A and K31A (Fig. 3C-D).

These findings suggest that RIL-seq chimera numbers do not fully capture the functional activity of Hfq, underscoring the need for further understanding how the pattern of chimera formation is affected by Hfq mutants. Given that R16A showed the most dramatic loss of S-chimeras, we performed additional analyses focusing on comparisons between WT and R16A.

### Chimera formation and R16A mutant defects are unaffected by the Flag tag

Recent studies have shown that the disordered C-terminus (CTD) of Hfq plays a role in regulating RNA binding and sRNA activity (8,29,44). In particular, combining either an R16A or a K31A mutation with deletion of the Hfq CTD had a synergistic effect on regulation (29). Thus, it seemed possible that the C-terminal Flag tag, used for capturing Hfq-RNA complexes, could be contributing to the loss of S-chimeras with R16A by effects on the CTD. To test whether the Flag tag affects S-chimera formation and exacerbates the defects observed in the R16A mutant, we adjusted the RIL-seq protocol to use a native Hfq antibody to isolate the Hfq-RNA complex instead of anti-Flag. This modification eliminated the need for a Flag tag at the C-terminus, while the other steps remained unchanged (Fig. 4A). Using antibodies against Hfq, we performed RIL-seq experiments with four strains (anti-Hfq): WT (Untagged), WT-Flag, R16A (Untagged), and R16A-Flag (Supplementary Table S6). The Circos plots and S-chimera numbers corresponding to these experiments are shown in Fig. 4B and 4C. In the WT-Flag strain, we observed an average of 4,339 S-chimeras (Fig. 4B and 4C), which is comparable to the 4,426 average S-chimeras obtained in anti-Flag RIL-seq (Fig. 1E). This demonstrates that the protocol using anti-Hfq antibody functioned well. In the WT Hfq strain (without the Flag tag), an average of 7,316 S-chimeras were identified (Fig. 4B and 4C), more than in the Flag-tagged strain (average of 4,339 S-chimeras). However, the similar patterns in the Circos plots between WT and WT-Flag suggest that the Flag tag does not markedly affect the distribution of S-chimeras formed. In parallel, the WT-Flag and R16A-Flag strains were also used for RIL-Seq, using the classic anti-FLAG protocol; this data has been deposited in GEO but not further analyzed here.

**Fig. 4.**
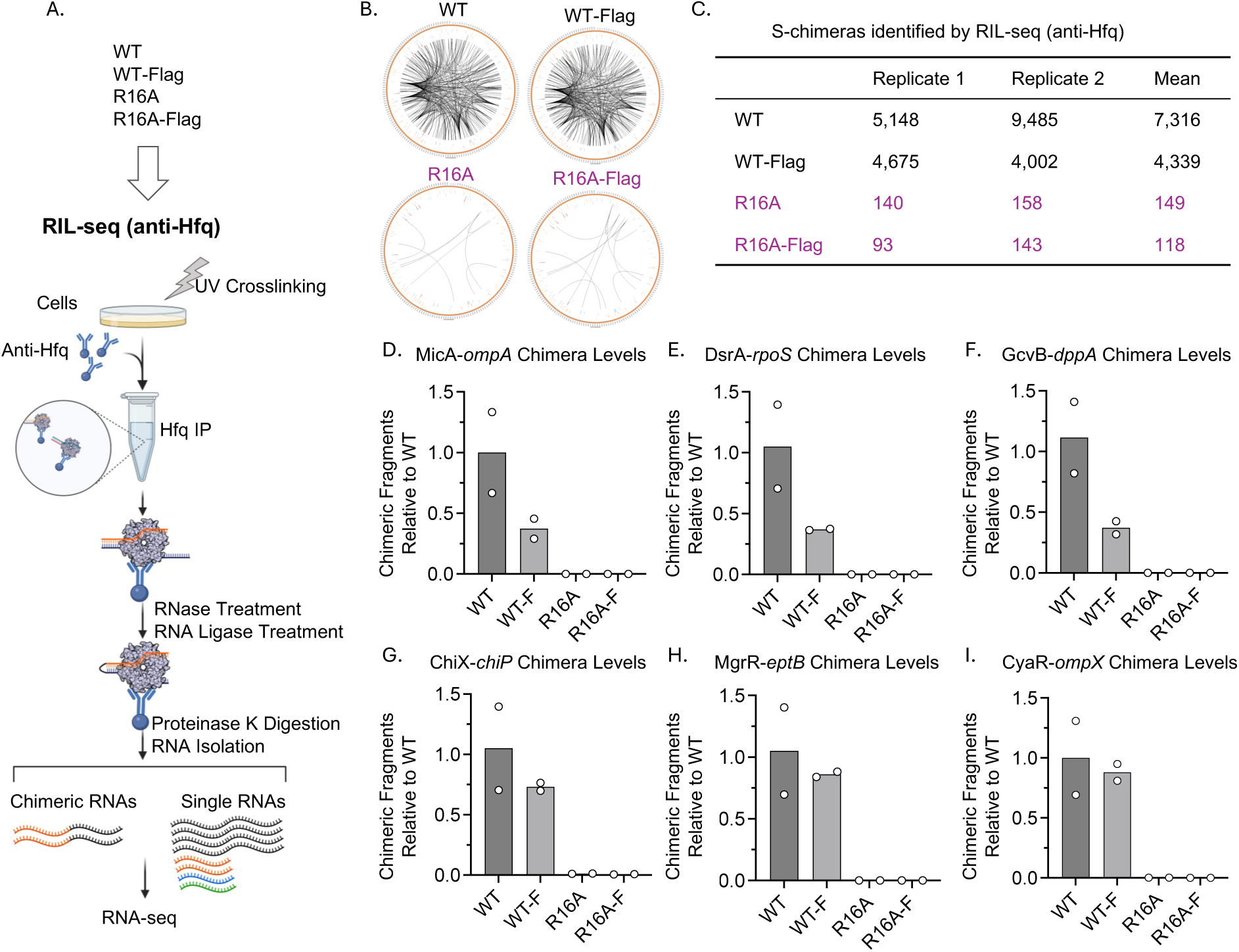
The Flag tag does not notably impact chimera formation in RIL-seq. (A) Workflow of RIL-seq (anti-Hfq) comparing WT and R16A Hfq with or without a Flag Tag. RIL-seq (anti-Hfq) was performed in WT (XL2001), WT-Flag (FDM1004), R16A (FDM1050), and R16A-Flag (FDM1006) to compare chimera formation across these strains. Hfq IP was conducted using an anti-Hfq antibody, while all other steps followed the same protocol as anti-Flag RIL-seq (Fig. 1A). sRNAs are shown in orange, mRNAs in black, and other Hfq-binding RNAs, such as tRNA and rRNA, are indicated in green and blue, respectively. This figure was created in BioRender (https://BioRender.com/jqqlsju). (B) Circos plots showing RNA interactomes in WT and R16A Hfq with or without a Flag tag. (C) Quantification of S-chimeras across conditions (Supplementary Table S6). Table displaying the number of RNA pairs with significant chimeric fragments identified by RIL-seq (anti-Hfq), with counts presented for two biological replicates and the calculated mean. (D–I) Chimera levels for representative sRNA-mRNA interactions, normalized to WT levels: (D) MicA-*ompA*, (E) DsrA-*rpoS*, (F) GcvB-*dppA*, (G) ChiX-*chiP*, (H) MgrR-*eptB* and (I) CyaR-*ompX*. Chimeric fragment levels were extracted from Supplementary Table S6, following the same approach as described for Figure 2B. Bars represent the average of two biological replicates, with individual replicate values shown as white dots.

For the R16A mutants, with or without the Flag tag, we identified a similarly low number of S-chimeras (Fig. 4B and 4C). The levels of chimeras for the six sRNA-mRNA pairs were also consistently low in the R16A strains for both Class I and Class II sRNAs (Fig. 4D-I). There was no significant difference in the number of chimeras for the six representative sRNA-mRNA pairs between the WT and WT-Flag strains. Overall, these data indicate that the R16A mutation disrupts Hfq-RNA interactions, which is not due to the combination of R16A with the Flag tag.

### The R16A mutation has distinct effects on Class I and Class II sRNA binding to Hfq

We next investigated whether S-chimeras were not detected because the R16A mutation disrupted RNA binding. Without effective RNA binding, Hfq cannot facilitate RNA pairing, leading to a reduction in chimeric RNA formation. In a previous study (21), R16A was found to destabilize Class I sRNAs and block the binding of the targets of Class II sRNAs, stabilizing the Class II sRNAs. Here, we revisited the Hfq interaction with sRNAs and their targets, based on both the RIL-seq procedure and a RIP-seq protocol (Fig. 5A), comparing RNA binding to WT Hfq and R16A, with and without the Flag tag.RIL-seq can provide a measure of general Hfq-bound RNAs by combining chimeric and single RNA fragments, with chimeras representing around 15% of all bound fragments in WT strains (see Supplementary Fig. S3). We re-analyzed the RIL-seq data, pooling together chimeras and single RNA fragments, mapping them to the genome, and calculating normalized Hfq IP RNA levels (Supplementary Table S7). Unlike RIL-seq (Fig. 4A), which was designed to capture RNA-RNA interactions, the RIP-seq protocol does not require cross-linking or *in vitro* RNase digestion, RNA ligation, and proteinase treatments (Fig. 5A). Bacterial cultures were grown to OD_600_ ∼1.0, rapidly collected by centrifugation, and lysed. Hfq IP was performed using an anti-Hfq antibody, and the co-purified RNA fraction was immediately isolated for cDNA library construction or Northern blot analysis. A similar analysis of WT compared to R16A IP was previously reported using RNA microarrays (23).

**Fig. 5.**
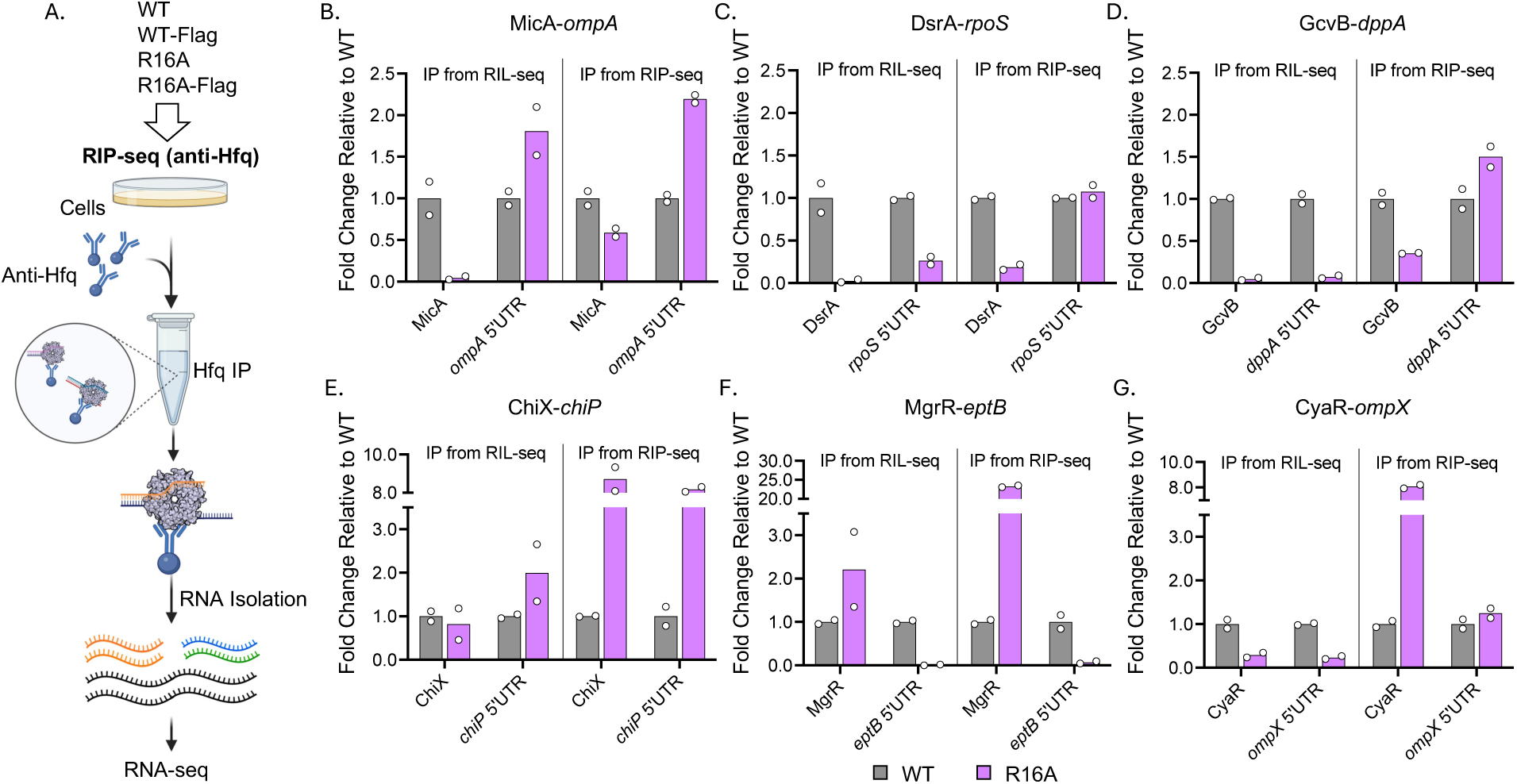
The R16A mutation differentially affects Class I and Class II sRNA binding to Hfq. (A) Workflow for RIP-seq conducted on WT (XL2001), WT-Flag (FDM1004), R16A (FDM1050), and R16A-Flag (FDM1006) strains. Immunoprecipitation was performed with anti-Hfq antibody, followed by RNA isolation and sequencing to identify RNA molecules bound by Hfq. sRNAs are shown in orange, mRNAs in black, and other Hfq-binding RNAs, such as tRNA and rRNA, are indicated in green and blue, respectively. This figure was created in BioRender (https://BioRender.com/jqqlsju). (B–G) RNA binding levels of sRNAs and their target mRNA interaction sites in R16A relative to WT, as determined by RIL-seq (left panels) and RIP-seq (right panels) for selected sRNA– mRNA pairs. IP levels were extracted from Supplementary Table S7 for RIL-seq, and Supplementary Table S8 for RIP-seq; relevant rows are highlighted in red in the Tables. Binding levels were normalized to WT to generate fold-change ratios. Each panel represents data for a specific sRNA or mRNA target. For most targets, the sRNA interaction region corresponds to the 5′UTR of the mRNA. Because the *rpoS* 5′UTR is not separately annotated due to its overlap with the upstream gene *nlpD*, we used the intergenic region between *rpoS* and *nlpD*, annotated as rpoS.nlpD.TU, to represent the DsrA binding site. Bars represent the average of two biological replicates, with individual replicate values shown as white dots.

As with the RIL-seq analysis, total RNA levels were determined in parallel with the RIP-seq experiments. The full RIP-seq data set, allowing comparison of Flag and non-Flag results, are in Supplementary Table S8. The Flag tags had very little effect on total RNA levels or RIP-seq (IP levels), as shown by PCA (Supplementary Fig. S5A) and volcano plots (Supplementary Fig. S5B,C and F,G). Comparisons of R16A to WT, with or without the Flag tag, showed modest differences in total RNA, but distinct differences in RIP-seq (Supplementary Fig. S5D,E and H, I), suggesting there is a defect in RNA binding to Hfq R16A.

We further compared the effect of the R16A mutation on RNA binding in the RIL-seq and RIP-seq protocols by examining the six selected sRNAs-target pairs (Fig. 5). The R16A mutant lost binding to the Class I sRNAs (Fig. 5B-D), consistent with previous work suggesting that this class of sRNAs directly interacts with the rim face of Hfq (21). However, the decrease in binding was more dramatic in the RIL-seq protocol (left panels) compared to RIP-seq (right panels), suggesting that the extra in vitro manipulation steps in RIL-seq may lead to further loss of these sRNAs. The reduced binding for R16A correlated with decreased total RNA levels of DsrA and GcvB (Supplementary Fig. S6B-C). These total RNA levels and Hfq IP levels of the sRNAs were further validated by northern analysis (Supplementary Fig. S7). Overall, the IP of the mRNAs was less affected, but again more RNA was retained by RIP-seq in R16A compared to RIL-seq (Fig. 5B-D).

Class II sRNAs were previously found to be more abundant and stable in the R16A mutant (21). This trend was observed in the RIP-seq results, where these sRNAs were 5 to 20-fold higher in the R16A mutant compared to WT (Fig. 5E-G, right panels) and in our northern analysis (Supplementary Fig. S7). In contrast, this increase was not seen with the RIL-seq protocol and varied with the sRNAs (Fig. 5E-G, left panels). The fate of the Class II mRNA targets also varied between the two protocols. Hfq capture of target mRNAs depends on both their abundance and their ability to bind Hfq. The results for the three tested targets here highlight that this balance can shift depending on the specific mRNA, and that RNA binding measured with the RIL-seq protocols is somewhat different from that measured with the more streamlined RIP-seq protocol.

Another approach to evaluating the extent of Hfq binding for each RNA is to calculate the enrichment ratio (IP/Total). A ratio greater than 1 indicates that most of the RNA is associated with Hfq, whereas a ratio below 1 suggests that a significant portion remains unbound. This measurement is useful for distinguishing between RNAs that are highly expressed but weakly bound to Hfq, where significant IP levels may still be observed but the enrichment ratio is very low, and those that are strongly associated with Hfq even if the RNA is expressed at a low level. We calculated enrichment ratios for the six model sRNAs and their mRNA targets using both RIL-seq (Supplementary Fig. S8A) and RIP-seq (Supplementary Fig. S8B) data. The overall trends were similar across both datasets. In the WT strain, all analyzed sRNAs were associated with Hfq, as expected, with enrichment ratios greater than 1. Among the target mRNAs, only *ompA* (an abundant mRNA) showed a low enrichment ratio, although this was also accompanied by a high adjusted *p*-value, suggesting limited reliability. In the R16A strain, the enrichment ratio and *p*-value were both poor for MicA in RIL-seq (Supplementary Fig. S8A) but were still strong in RIP-seq (Supplementary Fig. S8B). The mRNA targets of the Class II sRNAs MgrR and CyaR, *eptB* and *ompX* respectively, showed negative enrichment in both sets of data (Supplementary Fig. S8A-B). For *eptB*, this is due to a loss of binding in the R16A mutant (Fig. 5F), suggesting that its association with Hfq is particularly dependent on the R16 residue. In contrast, *ompX* exhibited a dramatic increase in total RNA levels in R16A (Supplementary Fig. S6F), presumably reflecting loss of sRNA negative regulation, while its IP level remained unchanged (Fig. 5G), leading to a reduced IP/Total ratio.

In summary, the rim face mutation in R16A reduces Class I sRNA binding, while Class II sRNAs are stabilized in this mutant, as previously found. However, the results here show different outcomes of the two protocols for capturing RNA on Hfq and also suggest that the RNA binding defects cannot account for the near-complete loss of S-chimeras from the R16A strain. We hypothesize that factors beyond RNA binding likely contribute to the failure to capture S-chimeras in the R16A strain by RIL-seq.

### Hfq R16A decreases the stability of sRNA-mRNA pairs on Hfq

A possible model to explain the reduced S-chimera detection in R16A, despite at least partial functionality, is that the sRNA-mRNA complex may dissociate from the R16A mutant more rapidly after pairing or more readily during post-IP processing. Since RIL-seq captures Hfq-bound chimeras, this dissociation would prevent the detection of these complexes. Such dissociation during the RIL-seq procedure is suggested by the higher loss of Class I sRNAs by RIL-seq compared to RIP-seq (Fig. 5).

To test if fewer post-IP steps results in the detection of more S-chimeras in R16A, we turned to the recently developed iRIL-seq (intracellular RIL-seq) (45) protocol. In the iRIL-seq protocol, RNA ligation occurs inside the cell rather than *in vitro*, minimizing post-IP steps. By inducing T4 RNA Ligase I from a plasmid, all spatially proximate RNA molecules—including interacting pairs—are ligated prior to Hfq IP. Following Hfq IP, RNA extraction is performed to isolate Hfq-associated RNA-RNA interactions; there is no cross-linking in this protocol, which resembles RIP-seq with the addition of the in vivo ligation step (Fig. 6A). iRIL-seq is derived from GRIL-seq (46) and Hi-GRIL-seq (47). In these methods, after T4 RNA Ligase I induction, total RNA is directly extracted and subjected to deep sequencing without Hfq IP (Fig. 6A). Unlike iRIL-seq, which focuses on Hfq-associated interactions, Hi-GRIL-seq is designed to capture global RNA-RNA interactions.

**Fig. 6.**
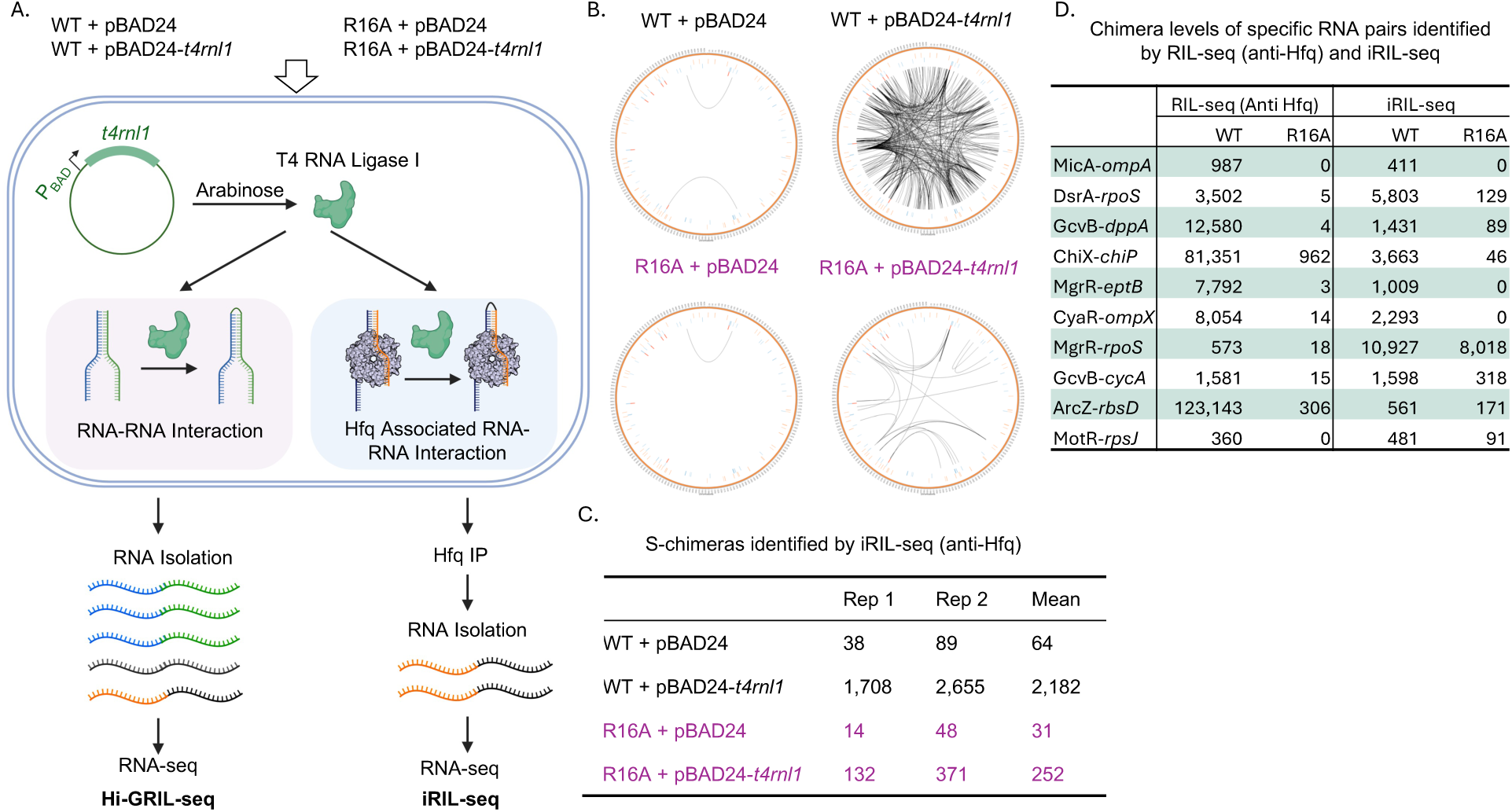
The R16A mutant retains more chimeras in iRIL-seq. (A) Schematic overview of Hi-GRIL-seq and iRIL-seq workflows. WT (XL2001) and the *hfq* R16A mutant (FDM1050) were transformed with either an empty vector (pBAD24) or a T4 RNA Ligase I-expressing vector (pBAD24-*t4rnl1*). Biological duplicates of each strain were cultured in LB (Lennox) with ampicillin at 37°C until reaching an OD600 of ∼0.5, at which point arabinose was added to induce enzyme expression. Cells were then grown to early stationary phase (OD600 ∼1.0), harvested, and processed for either Hfq IP followed by RNA isolation (iRIL-seq) or direct RNA extraction for total RNA sequencing (Hi-GRIL-seq). sRNAs are shown in orange, mRNAs in black, tRNA in green and rRNA in blue. This figure was created in BioRender (https://BioRender.com/jqqlsju). (B) Circos plots illustrating RNA interactomes in WT and R16A Hfq with either pBAD24 or pBAD24-*t4rnl1*, as detected by iRIL-seq. (C) Quantification of S-chimeras under different conditions (Supplementary Table S11). Table displaying the number of RNA pairs with significant chimeric fragments identified by iRIL-seq, with values shown for two biological replicates and their calculated mean. (D) Chimera levels of specific RNA pairs identified by RIL-seq (anti-Hfq) and iRIL-seq. The RNA pairs shown here include the six pairs analyzed in previous experiments as well as four additional pairs identified as having significant numbers of chimeras in the iRIL-seq R16A experiment. Chimera counts were extracted from Supplementary Table S6 (RIL-seq, anti-Hfq) and Supplementary Table S11 (iRIL-seq), using the same method described in Figure 2B. For each RNA pair, the values represent the mean from two biological replicates.

To assess how these different protocols affect Hfq-bound S-chimera detection, we performed iRIL-seq in four strains: WT Hfq carrying either an empty vector (pBAD24) or a T4 RNA Ligase I-expressing plasmid (pBAD24-*t4rnl1*), and the Hfq R16A mutant with the same plasmids. To evaluate total RNA levels and identify potential Hfq-independent S-chimeras, we also conducted Hi-GRIL-seq in parallel, using the same four strains as well as a Δ*hfq* strain carrying either pBAD24 or pBAD24-*t4rnl1*. Biological duplicates of each strain were grown in LB (Lennox) supplemented with ampicillin at 37°C until OD_600_ ∼ 0.5 (Supplementary Fig. S9A), at which point arabinose was added to induce enzyme expression (Supplementary Fig. S9A and S9B). Expression of T4 RNA Ligase I ultimately is toxic, as indicated by a subsequent reduction in growth rate (Supplementary Fig. S9A). Cells were harvested at the same OD_600_ as for RIL-seq (early stationary phase, OD_600_ ∼ 1.0), and processed for either Hfq IP followed by RNA isolation (iRIL-seq) or direct RNA extraction for total RNA sequencing (Hi-GRIL-seq) (Fig. 6A).

We first analyzed the total RNA and Hfq IP RNA levels from Hi-GRIL-seq and iRIL-seq, respectively (Supplementary Table S9). PCA analysis indicated a reasonable level of reproducibility of the duplicates (Supplementary Fig. S10A-B). Analysis of the six selected sRNAs and their target mRNAs for iRIL-seq (Supplementary Fig. S10C-H) showed that total RNA expression profiles were consistent with those observed for the previous RNA-seq (Supplementary Fig. S6) and Hfq binding profiles more closely resembled RIP-seq patterns than RIL-seq patterns (Fig. 5B-G, right panels, Supplementary Table S8). These findings suggest that iRIL-seq effectively captures Hfq-associated RNA profiles and further confirms that the more complex RIL-seq protocol affects the pattern of sRNAs captured by Hfq.

Next, we analyzed the chimeric RNAs obtained from Hi-GRIL-seq (Supplementary Table S10) and iRIL-seq (Supplementary Table S11). The background of chimeras captured in the absence of T4 RNA Ligase was 0.1% for Hi-GRIL-seq (Supplementary Fig. S11A). This increased to 0.6-0.7% chimeric RNAs for Hi-GRIL-seq in strains expressing the T4 RNA Ligase (Supplementary Fig. S11A), a level comparable to a previous study with *Pseudomonas aeruginosa* (47). The numbers of chimeric RNAs indicated intracellular RNA ligation was effective, but also suggested that only a small proportion of RNAs were either interacting or spatially proximate within the cell. One of the initial goals of performing Hi-GRIL-seq, particularly in the Δ*hfq* strain, was to determine if there were many Hfq-independent S-chimeras. However, relatively few S-chimeras were identified in the Δ*hfq* background (Supplementary Fig. S11B-C, Table S10) and those identified were enriched for highly abundant RNAs (tRNAs or rRNAs) that are presumably not due to association with Hfq (Supplementary Table S10). Therefore, our analysis focuses primarily on the iRIL-seq data.

For iRIL-seq, which enriches for Hfq-associated RNAs, chimeras represented 0.2% of total fragments in the absence of the T4 RNA Ligase; the number increased to 1.7% in the presence of the ligase (Supplementary Fig. S11D). This chimera fraction was less than in RIL-seq (anti-Flag) for WT (15%) (Supplementary Fig. S3). Further analysis of iRIL-seq chimeras revealed 2,183 S-chimeras in WT and 252 in R16A (Fig. 6B-C, Supplementary Table S11). While the ∼10-fold reduction in R16A is substantial, it is less severe than the ∼100-fold drop observed in RIL-Seq (anti-Flag) (Fig. 1D). This suggests that more S-chimeras were retained on HfqR16A in the iRIL-seq protocol.

iRIL-seq primarily captured interactions between mRNAs and sRNAs, although we also detected some chimeras involving abundant tRNAs and rRNAs (Supplementary Table S11). The top 25 S-chimeras for RIL-seq were compared to those for iRIL-seq and vice versa (Supplementary Table S12). For WT cells, almost all the top iRIL-seq S-chimeras were also found by RIL-seq; the exceptions involved tRNAs or rRNAs. About half of the top ranked RIL-seq S-chimeras were also found in iRIL-seq. Some S-chimeras not detected by iRIL-seq may be explained by the differences in growth conditions required for T4 RNA ligase expression, and a methodological bias of iRIL-seq, which cannot efficiently ligate primary transcripts bearing a 5′ triphosphate, thereby enriching for processed sRNAs (45).

As noted above, the number of iRIL-seq S-chimeras was reduced 10-fold in the R16A strain (Fig. 6C). About half of the top 25 most abundant S-chimeras for WT were not detected in either R16A dataset (Supplementary Table S13, top, marked as ND in column K or M). However, in contrast to the RIL-seq data (Supplementary Table S5), where a larger proportion of the top 20 S-chimeras from the R16A-Flag dataset was not observed in WT-Flag, 24 out of the top 25 most abundant iRIL-seq S-chimeras from R16A were also detected in WT (Supplementary Table S13, bottom). The one exception was a tRNA–rRNA S-chimera. Thus, in iRIL-seq the nature of the chimeras in R16A are more similar to the WT chimeras detected by both RIL-seq and iRIL-seq. This pattern, along with the 10-fold rather than 100-fold decrease in S-chimeras, suggests that the effect of the R16A mutation on Hfq-dependent binding and RNA interactions is disrupting pairing in a way that is much more sensitive to the RIL-seq capture protocol than to the iRIL-seq protocol.

This comparison of RIL-seq and iRIL-seq sensitivity to R16A was further examined using ten specific sRNA–mRNA pairs. The chimera counts for these RNA pairs in the WT and R16A strains, identified by RIL-seq (anti-Hfq) and iRIL-seq, are listed in Fig. 6D. Of the six representative sRNA–mRNA pairs analyzed previously, all were detected in the WT strain by iRIL-seq (Fig. 6D, top 6 RNA rows, Supplementary Table S11). As in RIL-seq (Fig. 4D–4I), iRIL-seq showed a significant reduction in chimeras for these pairs in R16A (Fig. 6D). However, iRIL-seq revealed certain sRNA-mRNA pairs that were undetectable or extremely low in R16A by RIL-seq but were reasonably abundant in R16A by iRIL-seq. *rpoS* is another known target of MgrR, with the interaction of this sRNA with the *rpoS* 5’ UTR previously detected using both iRIL-seq and another GRIL-seq related method (45),(48); RpoS levels were also shown to be affected by MgrR overexpression in *E. coli* (49). In our study, the MgrR–*rpoS* pair showed the highest chimera levels by iRIL-seq in both WT and R16A strains (Supplementary Table S13). In contrast, this interaction was much less prominent in RIL-seq, ranking below 200 in WT (Supplementary Table S6) and being barely detectable in R16A (Fig. 6D). Additionally, *cycA*, another target of GcvB (50), showed over 20-fold more GcvB–*cycA* chimeras in R16A by iRIL-seq than by RIL-seq (318 vs. 15), while both methods detected similar levels in the WT strain (Fig. 6D). Additional examples included ArcZ–*rbsD*, a regulatory pair in which *rbsD* mRNA acts as a decoy RNA, negatively regulating ArcZ, an activator of *rpoS* translation (51). The ArcZ-*rbsD* chimeras were abundant in WT in both RIL-seq (Supplementary Table S4 and S6) and iRIL-seq (Supplementary Table S11). While this interaction was barely detected in R16A by RIL-seq, R16A retained approximately 30% of the chimera level observed in WT in the iRIL-seq data (Fig. 6D). Similarly, the interaction between MotR, a 5′UTR-derived sRNA, and *rpsJ* (52), the first gene of the S10 operon, was recovered exclusively by iRIL-seq for R16A (Fig. 6D).

Overall, these results are consistent with Hfq R16A still able to facilitate sRNA-mRNA interactions. However, the sRNA-mRNA pairs appear to be less stably bound to R16A compared to WT Hfq. Additionally, the different distributions of sRNA-mRNA S-chimeras for WT Hfq versus R16A in the RIL-seq and iRIL-seq data may provide an indication of features of individual sRNA-mRNAs pairs that impact continued binding to Hfq after base pairing.

## DISCUSSION

The recent development of methods for capturing sRNAs and their targets on RNA binding proteins (RIL-seq, RIP-seq, and iRIL-seq) has greatly expanded our understanding of sRNA-dependent regulation in bacteria (24,25,27,45–47). RIL-seq, in which RNAs bound together to Hfq are ligated, leads to the identification of S-chimeras that reflect known sRNA-mRNA pairing. We began this study to determine how mutations in the critical residues on the RNA binding faces of Hfq affected formation of these S-chimeras, with the expectation that this would help elucidate the roles of these residues during *in vivo* pairing. In the RIL-seq analysis, the proximal face D9A mutant and the distal face Y25D did not notably change the number of S-chimeras; the distal face K31A mutant reduced the number of S-chimeras by almost 4-fold. However, the rim face mutant R16A showed a near-complete loss of S-chimeras. This result suggested that the rim face had a uniquely important role in capturing RNA-RNA interactions by the RIL-seq protocol. This unexpected and dramatic role of the R16A rim mutant led us to further investigate the basis for this loss of S-chimeras.

Because RIL-seq has previously been carried out using C-terminally Flag-tagged Hfq, we first investigated the role of the tag on Hfq in S-chimera formation, in both WT and R16A cells, and found very little effect of this tag on total gene expression (Total RNA-seq, Supplementary Fig. S2C, Table S3, Fig. S5B-C, Table S8), RNA binding ability (RIP-seq, Supplementary Fig. S5F-G, Table S8), or S-chimera formation (RIL-seq (anti-Hfq), Fig. 4 and Supplementary Table S6). Certainly, the Flag tag did not explain the lack of S-chimeras in R16A mutants in the original RIL-seq analysis. We were able to demonstrate that the RIL-seq protocol works well using antibodies to Hfq, an advantage in cases where a tag may disrupt protein function or the genetics to integrate a tag in vivo are not available. For Hfq, the Flag tag had a minimal impact on Hfq function and thus remains a reliable and valuable tool for experimental studies, enabling efficient detection and immunoprecipitation while preserving most RNA interactions.

### Role of Hfq R16 in RNA Binding and sRNA-mRNA Pairing

Mutations in the Hfq rim, including the R16A mutant, have been shown to reduce both the binding and stability of Class I sRNAs, leading to lower levels of these sRNAs (21), also supported by our RIP-seq analysis. This reduction in sRNA levels was not sufficient to abolish regulatory activity, as previously observed (22) and supported here by our observations of total RNA levels as well as RpoS levels in the R16A strain. This makes the near-complete loss of S-chimeras for Class I sRNAs in RIL-seq particularly surprising. Class II sRNAs, in contrast, exhibit increased abundance in the R16A mutant (21). Given that the targets of Class II sRNAs bind the Hfq rim face, it may not be surprising to observe fewer S-chimeras forming for these sRNAs in the R16A background. We note that the targets of Class I sRNAs bind the distal face, and yet neither of the two distal face mutants, K31A or Y25D, dramatically blocked chimeras with these sRNAs. This suggests an additional role for R16A. Interestingly, *in vitro* studies have shown that Hfq rim mutants, including R16A, do not completely prevent RNA binding. Instead, these mutations have a notable and unique impact on the RNA annealing activity of Hfq, impairing its ability to facilitate sRNA-mRNA duplex formation (19). Collectively, these results suggest that the selective defect in S-chimeras in R16A may not primarily reflect a rim function in RNA recruitment but rather a more prominent role in facilitating or stabilizing sRNA-mRNA pairing (Fig. 7).

**Fig. 7.**
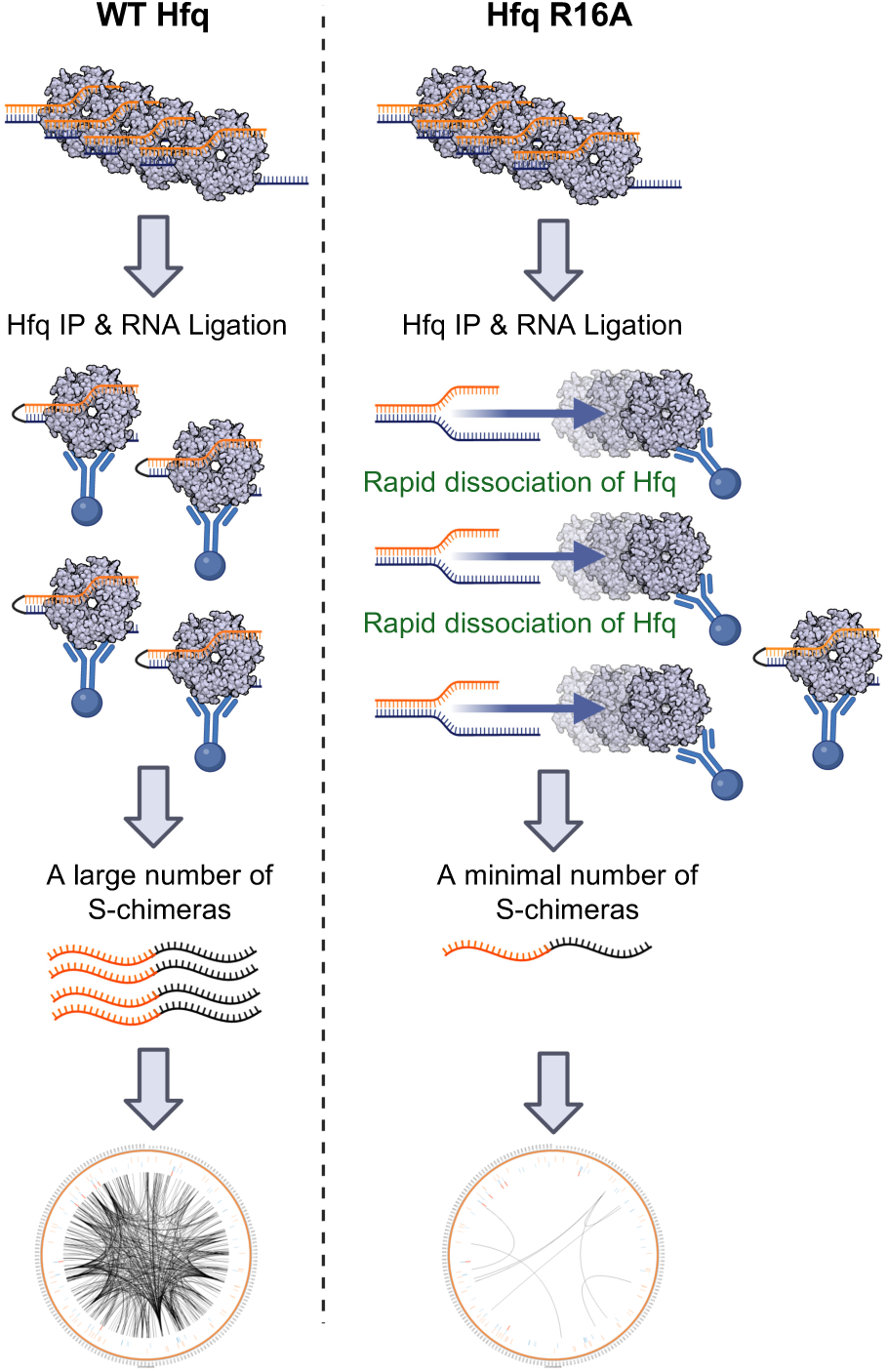
The R16A mutant reduces the stability of sRNA-mRNA pairs on Hfq. On WT Hfq (left), sRNA-mRNA pairs are efficiently stabilized, allowing for the formation of chimeric RNAs that can be captured and sequenced, resulting in a dense RNA interactome. In contrast, in the Hfq R16A mutant (right), the mutation disrupts the rim face, leading to rapid dissociation of sRNA-mRNA pairs. This instability results in substantially fewer chimeric RNAs being detected by RIL-seq, as many transient interactions are lost before sequencing. The Circos plots at the bottom illustrate the difference in detected RNA interactions between WT and R16A. This figure was created in BioRender (https://BioRender.com/jqqlsju)

We employed the iRIL-seq approach to test whether capturing chimeras *in vivo* (by expressing T4 RNA ligase during cell growth) could bypass the S-chimera formation defect observed in RIL-seq (where chimeras are generated after RNA crosslinked to Hfq is isolated from the cell). In our experiments, iRIL-seq revealed a 10-fold reduction in S-chimeras for R16A compared to WT, in contrast to the 100-fold decrease observed using RIL-seq. We interpret this finding as evidence that sRNA and mRNA binding to Hfq is not completely abolished by the R16A mutation—allowing for some regulatory function and detection of S-chimeras by iRIL-seq. Presumably, the interaction is likely destabilized to a degree that prevents effective chimera formation during the more stringent post-IP processing steps required in the RIL-seq protocol.

### Strengths and Limitations of RIL-seq and iRIL-seq

Both RIL-seq and iRIL-seq are powerful methods for defining RNA-RNA interactions *in vivo*, each with distinct advantages. RIL-seq is a well-established protocol that excels at capturing stable Hfq-mediated interactions, enabling high-confidence identification of chimeric RNAs. A key benefit of RIL-seq is that it does not require antibiotics or inducers like arabinose, which helps maintain natural cellular conditions, avoiding possible physiological effects on chimera formation during changes in growth. However, the protocol includes multiple post-immunoprecipitation *in vitro* processing steps, such as RNase digestion and RNA ligation, which may lead to the loss of transient or less stable interactions.

In contrast, iRIL-seq minimizes post-IP processing by performing RNA ligation inside the cell, which allows for better retention of transient RNA interactions that may dissociate during standard RIL-seq steps. This makes iRIL-seq particularly useful for detecting short-lived or dynamic RNA pairings. Additionally, iRIL-seq is simpler and faster to perform than standard RIL-seq, making it attractive for comparing mutants or different growth conditions. However, iRIL-seq requires expression of T4 RNA ligase I, which involves plasmid maintenance, antibiotic selection, and induction with arabinose. These requirements can introduce physiological stress, as evidenced by the growth defects observed in strains carrying the enzyme-expressing plasmid. Moreover, because T4 RNA ligase I can only ligate RNAs with a 5′ monophosphate, iRIL-seq may fail to capture primary transcripts that retain a 5′ triphosphate, potentially excluding some sRNAs from detection (45).

The differences between these protocols is exemplified by the significant differences we observed in the S-chimera profiles generated by RIL-seq and iRIL-seq. Certain S-chimeras, such as the highly abundant FnrS–*ytfK* pair detected by RIL-seq, were not observed in iRIL-seq. Conversely, the most abundant S-chimera identified by iRIL-seq, MgrR–*nlpD* (*rpoS*), was ranked low in the RIL-seq dataset. These discrepancies may reflect the distinct growth conditions and expression systems required by the two protocols, or possibly other important aspects of chimera formation worth future study.

Overall, RIL-seq is highly effective for identifying stable, native RNA interactions, while iRIL-seq offers an alternative route to detecting transient or weaker interactions that might be missed using standard methods. Depending on the research objective, the two approaches can serve as complementary strategies to yield a more complete view of RNA-RNA regulatory networks and provide further insights into nuanced features of RNA-protein interactions.

## Supporting information

Supplemental Protocol File 4

Supplemental Protocol File 2

Supplemental Protocol File 1

Supplemental Protocol File 3

## ACKNOWLEDGEMENTS

We thank Nadim Majdalani for the gift strain NM572. We thank Sarah Svensson, Sarah Woodson and members of Gottesman lab for critical reading and comments. We also thank that NICHD Molecular Genomics Core, particularly T. Li, for library sequencing. This work utilized the computational resources of the NIH HPC Biowulf cluster (http://hpc.nih.gov).

S.G. and G.S. conceived this project. X.L., A.Z., F.D.M. and P.P.A. performed the experiments. C.E. and R.K.D. processed the RNA-seq data. X.L., A.Z., C.E., R.K.D., G.S. and S.G. analyzed the data and interpreted the results. X.L. wrote the first draft and edited the paper. P.P.A., G.S. and S.G. reviewed and edited the paper.

## SUPPLEMENTARY DATA

Supplementary Data are available at NAR online; titles of all Supplementary figures and tables can be found in the Supplementary Information file.

## CONFLICT OF INTEREST

None declared.

## FUNDING

This work was funded by the National Institutes of Health, Division of Intramural Research: the Center for Cancer Research of the National Cancer Institute [ZIA BC 008714 to S.G.]; the *Eunice Kennedy Shriver* National Institute of Child Health and Human Development [1ZIAHD01608 to G.S.], [1ZICHD008986 to R.K.D], and [1ZIAHD008995 to P.P.A.]; the National Institute of Allergy and Infectious Diseases [1ZIAAI001395 to P.P.A.].

## DATA AVAILABILITY

The data here can be visualized at the UCSC browser with the following links. The four links represent two sets of replicates, as follows:

RIL-seq (anti-Flag) rep 1 and rep 2: Data from Supplementary Tables S3 and S4; relevant to Figures 1 and 2:

https://www.nichd.nih.gov/about/org/dir/other-facilities/cores/bioinformatics/data/luo-RILSeq_antiFlag_rep1

https://www.nichd.nih.gov/about/org/dir/other-facilities/cores/bioinformatics/data/luo-RILSeq_antiFlag_rep2

RIL-seq anti-Hfq and iRIL-seq rep 1 and rep 2: Data from RIL-seq, Supplementary Tables S6 and S7 and iRIL-seq Supplementary Table S9 and S11and Figures 4-6.

https://www.nichd.nih.gov/about/org/dir/other-facilities/cores/bioinformatics/data/luo-RILSeq_antiHfq_and_iRILseq_rep1

https://www.nichd.nih.gov/about/org/dir/other-facilities/cores/bioinformatics/data/luo-RILSeq_antiHfq_and_iRILseq_rep2

The raw sequencing data discussed in this study is accessible in GEO under the accessions GSE296794 (https://www.ncbi.nlm.nih.gov/geo/query/acc.cgi?acc=GSE296794) for the Total RNA-seq and RIP-seq experiments, GSE296795 (https://www.ncbi.nlm.nih.gov/geo/query/acc.cgi?acc=GSE296795) for all the RIL-seq experiments (both anti-Flag and anti-Hfq), and GSE296796 (https://www.ncbi.nlm.nih.gov/geo/query/acc.cgi?acc=GSE296796) for Hi-GRIL-seq and iRIL-seq experiments.

## Notes

### Competing Interest Statement

The authors have declared no competing interest.

