## Supplemental Protocol File 2 for "RNA-RNA Interactome Approaches Provide *in vivo* Evidence for a Critical Role of the Hfq Rim Face in sRNA-mRNA Pairing"

**Protocol for RNA-seq (with DASH)**

This protocol details the steps for cDNA library construction. In the protocol described here, rRNA is depleated using DASH.

This protocol was adapted and optimized based on the following previously published methods:

- cDNA library construction was adapted from the RIL-seq protocol described in PMID: 29215635.
- rRNA depletion was performed using DASH (Depletion of Abundant Sequences by Hybridization), modified from PMID: 32345633.
- Barcode sequences are described in PMID: 25730492 (where additional sequences can be found)

_________________________________________________________________________________

**DAY 0**

**Set up the night before for total RNA samples isolated previously**

1| Analyze the RNA samples (diluted 1/10) using RNA ScreenTape in the TapeStation to assess RNA quantity and size distribution.

2| Measure the RNA concentration using a Nanodrop and calculate up to 400 ng of RNA and complete the volume to 15 μL with DEPC water in PCR tubes. Add 1 μL of recombinant RNase inhibitor (40 U/µL).

**POSSIBLE PAUSE:** Continue with RNA-seq library construction or store the samples at −80 °C for up to 1 month.

_________________________________________________________________________________

**DAY 1**

### Fragmentation and DNase–FastAP combined treatment (45 min)

**CRITICAL:** RNA-seq libraries are constructed using the RNAtag-Seq protocol with a few modifications to allow capture of short RNA fragments. The library construction process includes a step of random RNA fragmentation. This step is required to obtain full coverage of long RNAs.

3| Add 4 μL of 10X FastAP buffer (Thermo Fisher Scientific, cat. no. EF0654) to the RNA tubes from Step 2 (total reaction volume is 20 μL), mix well and incubate the tubes in a preheated thermal cycler for 3 min at 92 °C. Remove the tubes quickly, put them in and out of ice in the same order.

(If RNA is partially degraded (RIN<7), fragment for 3 min or less at 92 °C – prevents over-fragmenting samples)

4| Place the samples on ice and prepare DNase–phosphatase reaction mix (20 μL per sample;
*x* samples + 3 extra samples):

| **Reagent** | **Volume (µL) per reaction** | **Final concentration (volume = 40 µL)** | **Per 12 samples = master mix for 15 samples (µL)** |
| --- | --- | --- | --- |
| Recombinant RNase inhibitor (40 U/µL) | 1 | 1 U/µL | 15 |
| TURBO DNase (2 U/µL) | 4 | 0.2 U/µL | 60 |
| FastAP (1 U/µL) | 10 | 0.25 U/µL | 150 |
| DEPC H_2_O | 5 |  | 75 |
| Mix total volume | 20 |  | 300 |

5| Add 20 µL of the mixture to each tube containing fragmented RNA, mix well and then incubate the tubes in a thermal cycler for 30 min at 37 °C.

### RNA cleanup using RNA Clean & Concentrator-5 kit (1 h)

6| Increase the reaction volume to 80 μL by adding 40 μL of nuclease-free water and transfer the mixture to a 1.5-mL LoBind tube (Eppendorf, cat. no. 022431021). Clean up the RNA according to the Zymo kit standard protocol for recovering RNA species of >17 nt. In the final step, elute the RNA with 8 μL of nuclease-free water and put on ice.

RNA cleanup using RNA Clean & Concentrator- 5 kit (Zymo Research, cat. no. R1014):

a. Add 160 µL of RNA Binding Buffer to each sample and mix.
b. Add 240 µL of 100% ethanol and mix.
c. Transfer the sample to the Zymo-Spin IC Column in a Collection Tube and centrifuge for 30 sec (13,000 rcf= 11,766 rpm, RT). Discard the flow-through.
d. Add 400 µL of RNA Prep Buffer to the column and centrifuge for 30 sec. Discard the flow-through.
e. Add 700 µL of RNA Wash buffer to the column and centrifuge for 30 sec. Discard the flow-through.
f. Add 400 µL of RNA Wash buffer to the column and centrifuge for 2 min to ensure complete removal of the wash buffer. Transfer the column carefully into an RNase- free tube (not provided)
g. Add 8 µL of DNase/RNase-Free Water directly to the column matrix, sit 1 min and centrifuge for 30 sec.

7| Optional: As a quality control, use an HS-RNA ScreenTape in the Agilent TapeStation (or use BioAnalyzer) to analyze random samples and check their fragmentation profile.

### Ligation of a 3’ adaptor (RNA/DNA) (2.5 h)

8| Make a list of the barcoded adaptors and the corresponding samples that will be ligated. The barcoded adaptors are listed in Table 1 of PMID: 29215635 and more in PMID: 25730492. In a PCR tube, mix 5 μL of dephosphorylated RNA from Step 6 with 1 μL of the respective barcoded adaptor (100 μM). Heat the tube at 70 °C for 2 min and place it on ice.

9| Set up the following ligation mix (14 μL per sample; *x* samples + 3 extra samples) at room temperature.

| **Reagent** | **Volume (µL) per reaction** | **Final concentration**  **(volume = 20 µL)** | **Per 12 samples = master mix for 15 samples (µL)** |
| --- | --- | --- | --- |
| T4 RNA ligase buffer, 10X | 2 |  | 30 |
| DMSO (100% (vol/vol)) | 1.8 |  | 27 |
| ATP (100 mM) | 0.2 | 1 mM | 3 |
| PEG 8000 (50% (wt/vol)) | 8 |  | 120 |
| Recombinant RNase inhibitor (40 U/µL) | 0.3 | 0.6 U/µL | 4.5 |
| T4 RNA ligase 1 (30,000 U/µL) | 1.7 | 2.55 U/µL | 25.5 |
| Mix total volume | 14 |  | 210 |

**CRITICAL:** Set up the mixture at room temperature to prevent DMSO precipitation. Pipette very slowly for accurate aspiration of PEG, as it is very viscous. Mix well by tapping the tube, as the solution is very viscous, and spin down.

10| Add 14 μL of ligation mix to each tube containing 6 μL of denatured RNA + adaptor. Mix well by tapping the tubes, as the solution is very viscous, and incubate the tubes at 22 °C for 2 h.

### Pooling the libraries and RNA cleanup using RNA Clean & Concentrator-5 kit (30 min)

11| Add 60 μL of RLT buffer (Qiagen, cat. no. 79216) to each sample to inhibit ligase activity and mix the solution well.

12| Pool three samples together into one tube by mixing 80 µL (total amount) of samples 1-3, 4-6, 7-9 and 10-12, ending with 4 tubes with 240 µL each. Then clean up using the RNA Clean & Concentrator-5 kit (Zymo Research, cat. no. R1014). Follow the kit’s standard protocol for recovering RNA species >17 nt.

RNA cleanup using RNA Clean & Concentrator- 5 kit (second clean-up):

a. Add 480 µL of RNA Binding Buffer to each sample and mix.
b. Add 720 µL of 100% ethanol and mix.
c. Transfer samples to two Zymo-Spin IC Column in a Collection Tube (ex. Combine samples 1-6 in one column and 7-12 in the other) and centrifuge for 30 sec. Discard the flow-through. Repeat the process using the same columns until all samples have been processed – 700 µL at a time.
d. Add 400 µL of RNA Prep Buffer to the column and centrifuge for 30 sec. Discard the flow-through.
e. Add 700 µL of RNA Wash Buffer to the column and centrifuge for 30 sec. Discard the flow-through.
f. Add 400 µL of RNA Wash Buffer to the column and centrifuge for 2 min to ensure complete removal of the wash buffer. Transfer the column carefully into an RNase-free tube (not provided).
g. Add 6.5 µL of DNase/RNase-Free Water directly to the column matrix and centrifuge for 30 sec.
h. Repeat: Add the elute back to the same column matrix and centrifuge for 30 sec.
i. Combine two tubes, so total volume = 6.5 x 2= 13 µL.
j. Store at -80°C.

**CRITICAL:** If you have more than eight samples, load the mixture of pooled samples, binding buffer and ethanol onto two columns. Combine the eluates in one tube, final volume = 13 μL.

_________________________________________________________________________________

**DAY 2**

**First-strand cDNA synthesis using the superscript III First-strand synthesis system (1.25 h)**

13| Add 11 µL of the eluted sample from Step 12 to a new PCR tube and add 1 µL of 50 µM AR2 primer (sequence: 5’-TACACGACGCTCTTCCGAT-3’; also refer to Table 1 of PMID: 29215635). Mix well.

14| Heat the mixture to 70 °C for 2 min and immediately place it on ice.

15| Set up the following reverse transcription mix on ice:

| **Reagent** | **Volume (µL) per reaction** | **Final concentration (25 µL)** |
| --- | --- | --- |
| dNTP mix (10 mM) | 1.25 | 0.5 mM |
| 10× RT buffer | 2.5 |  |
| MgCl2 (25 mM) | 5 | 5 mM |
| DTT (0.1 M) | 2.5 | 0.01 M |
| RNaseOUT (40 U/µL) | 0.5 | 0.8 U/µL |
| SuperScript III RT (200 U/µL) | 1.25 | 10 U/µL |
| Mix total volume | 13 |  |

16| Add 13 µL of reverse transcription mix to the mixture from Step 14. Mix well and incubate the mixture at 50 °C for 55 min.

**POSSIBLE PAUSE:** Store the sample at −80°C for up to 1 month or proceed to the next step.

### RNA degradation (20 min)

17| Add 2.5 µL of 1N NaOH (2 µL of 5N NaOH and 8 µL DEPC H_2_O) to the tube from Step 16 and incubate it at 70 °C for 12 min.

18| Add 5 µL of freshly diluted 0.5 M acetic acid (1/33 from original stock (16.5 M from Sigma) – 30 µL of stock + 970 µL of H_2_O) and mix well.

### cDNA cleanup (45 min)

19| Add 7.5 µL of nuclease-free water for a final volume of 40 µL and transfer it to a new Eppendorf tube.

20| Add 1.5X (60 µL) isopropanol and 2.5X (100 µL) RNAclean XP beads (Beckman Coulter, cat. no. A63987), mix by pipetting 15X and incubate the tube at room temperature for 15 min.

21| Place the tube on a magnetic rack for about 5 min until the solution is clear and discard the supernatant.

22| Wash the beads with 200 µL of freshly prepared 80% (vol/vol) ethanol without removing the tube from the magnetic rack. Incubate the tube for 30 s and discard the supernatant. Repeat the wash once more, and discard all the remaining ethanol droplets.

23| Leave the tube open on the magnetic rack and allow the beads to air-dry at room temperature for 10 min.

24| Remove the tube from the magnetic rack, resuspend the beads in 5 µL of nuclease-free water and incubate the tube for 2 min at room temperature.

**CRITICAL:** Keep the cDNA in the tube with the beads for the following ligation step.

### Ligation of a second adaptor at the cDNA 3’ end (ssDNA/ssDNA) (overnight)

25| Transfer DNA with beads to a new PCR tube and add 2 µL of 40 µM 3Tr3 adaptor (sequence: P-AGATCGGAAGAGCACACGTCTG-ddC; also refer to Table 1 of PMID: 29215635) to the cDNA+beads solution, incubate the tube at 75°C for 3 min and place it on ice.

26| Set up the following ligation mix at room temperature:

| **Reagent** | **Volume (µL) per reaction** | **Final concentration (20 µL)** |
| --- | --- | --- |
| T4 RNA ligase buffer, 10× | 2 |  |
| DMSO (100% (vol/vol)) | 0.8 |  |
| ATP (100 mM) | 0.2 | 1 mM |
| PEG 8000 (50% (wt/vol)) | 8.5 |  |
| T4 RNA ligase 1 (30,000 U/mL) | 1.5 | 2.25 U/mL |
| Mix total volume | 13 |  |

**CRITICAL:** Set up the mix at room temperature to prevent DMSO precipitation. Pipette very slowly for accurate aspiration of PEG, as it is very viscous. Mix well by tapping the tube, as the mixture is very viscous, and spin down.

27| Add 13 µL of ligation mix to the tube from Step 25. Mix well by tapping the tubes, as the solution is very viscous. Spin down and incubate the tube overnight at 22°C.

_________________________________________________________________________________

**DAY 3**

### Cleanup of cDNA (45 min)

28| Increase the volume to 40 µL by adding 20 µL of nuclease-free water and transfer to a new Eppendorf tube. Add 1.5X (60 µL) isopropanol and 2.5X (100 µL) RNAClean XP beads, mix by pipetting 15X and incubate the tube at room temperature for 15 min.

29| Place the tube on a magnetic rack for ~5 min until the solution is clear and discard the supernatant.

30| Wash the beads with 200 µL of freshly prepared 80% (vol/vol) ethanol (800 µL EtOH+ 200 µL DEPC H_2_O) without removing the tube from the magnetic rack. Incubate the tube for 30 s and discard the supernatant. Repeat the wash once more and discard all the remaining ethanol droplets. Leave the tube open to air-dry for 10 min.

31| Resuspend the dried beads in 25 µL of nuclease-free water. Incubate the tube for 2 min at room temperature and then place the tube on the magnetic rack. **SAVE LIQUID** After the solution appears clear (1 min), transfer the supernatant containing the eluted cDNA to a new Eppendorf tube.

**POSSIBLE PAUSE:** Store the sample at −80 °C for up to 1 month or proceed to the next step.

### 2nd cleanup of cDNA to remove the remaining adaptors (45 min)

32| Add 1.5X (37.5 µL) isopropanol and 2.5X (62.5 µL) RNAClean XP beads, mix by pipetting 15X and incubate the tube at room temperature for 15 min.

33| Place the tube on a magnetic rack for ~5 min until the solution is clear and discard the supernatant.

34| Wash the beads with 200 µL of freshly prepared 80% (vol/vol) ethanol without removing the tube from the magnetic rack. Incubate the tube for 30 s and discard the supernatant. Repeat the wash once more and discard all the remaining ethanol droplets. Leave the tube open to air-dry for 5 min.

35| Resuspend the dried beads in 18 µL of nuclease-free water. Incubate the tube for 2 min at room temperature and then place the tube on the magnetic rack. **SAVE LIQUID** After the solution appears clear (1 min), transfer the supernatant containing the eluted cDNA to a new Eppendorf tube.

**POSSIBLE PAUSE:** Store the sample at −80 °C for up to 5 years or proceed to the next step.

**DASH pre-amplification PCR (30 min)**

36| Prepare the following PCR mix:

| **Reagent** | **Volume (µL) per reaction** |
| --- | --- |
| Nuclease-free water | 0.5 |
| P5_Enr primer (12.5 µM) | 2 |
| P7_BC1_Enr primer (12.5 µM) | 2 |
| cDNA | 8 |
| KAPA HIFI HotStart ReadyMix, 2X | 12.5 |
| Mix total volume | 25 |

37| Use the total 25 µL of PCR mix in one PCR tube.

38| Amplify the cDNA using 2 cycles, with the following PCR program:

| **Step** | **Temperature (˚C)** | **Time** | **Number of cycles** |
| --- | --- | --- | --- |
| Initial denaturation | 95 | 3 min | 1 |
| Denaturation | 95 | 30 s |  |
| Annealing | 55 | 30 s | 2 |
| Extension | 72 | 30 s |  |
| Final extension | 72 | 10 min | 1 |
| Hold | 4 | ∞ |  |

**POSSIBLE PAUSE:** Store the amplified libraries at −80 °C for up to 1 month or proceed to the next step.

### PCR cleanup (45 min)

39| Transfer 25 µL PCR product into a new Eppendorf tube, add 1.4X RNAclean XP beads (35 µL), mix by pipetting 15X and incubate the tube at room temperature for 15 min.

40| Place the tube on a magnetic rack for about 5 min until solution is clear and discard the supernatant.

41| Wash the beads with 200 µL of freshly prepared 80% (vol/vol) ethanol without removing the tube from the magnetic rack. Incubate the tube for 30 s and discard the supernatant. Repeat the wash once more and discard all the remaining ethanol droplets. Leave the tube open to air-dry for 5 min.

42| Resuspend the dried beads in 11 µL of nuclease-free water. Incubate the tube for 2 min at room temperature and then place the tube on the magnetic rack. **SAVE LIQUID** After the solution appears clear, transfer the supernatant containing the eluted PCR fragments to a new Eppendorf tube.

43| Loading 1 µL of the PCR product on a NanoDrop to measure its concentration; the typical concentration is 10–20 ng/µL. NanoDrop the sgRNA pool to determine the concentration and perform Agilent TapeStation (or BioAnalyzer) analysis to determine the size of the sgRNA pool. Note the sgRNA pool is *in vitro* transcribed for a specific bacterial strain, exactly as described in PMID: 32345633.

**rRNA depletion using DASH (2.5 h)**

44| Use the DASH calculation table (Protocol_File_4_DASH_calculator_for E. coli _rRNA_depletion) to determine the amount of Cas9 and sgRNA pool to use for the DASH reaction. Add the cDNA concentration, sgRNA concentration, and sgRNA average size into calculator – yellow highlight. Example:

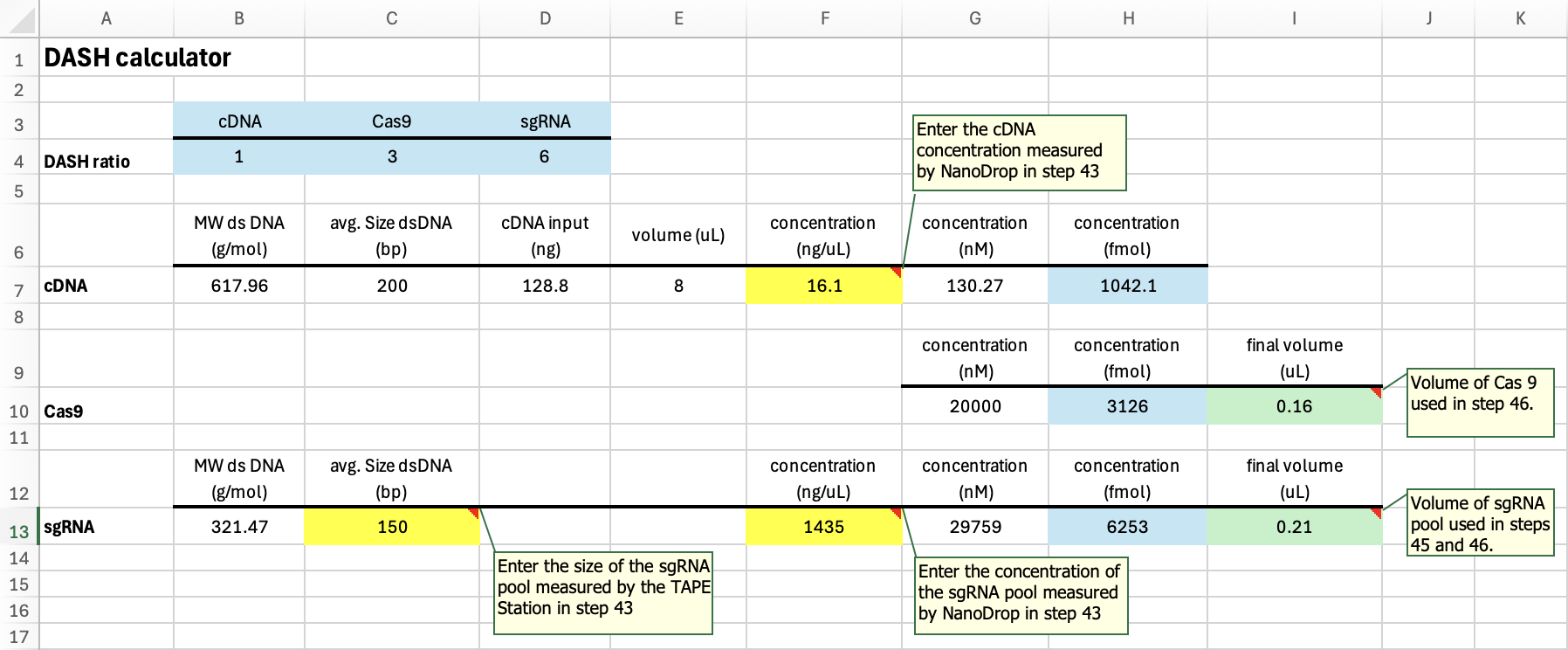

45| Add calculated sgRNA (ex. 0.21 μL) and nuclease-free water (amount per the table below) to a new PCR tube, incubate the tube at 94°C for 3 min, 4°C for 5 min.

46| Prepare the DASH reaction:

| **Reagent** | **Volume (****µL ) per reaction** | **Example reaction (µL)** |
| --- | --- | --- |
| sgRNA pool | per the calculation | **0.21** |
| Nuclease-free water | up to 10.5 | 8.28 |
| Cas9 buffer (3.1 NEB buffer) 10x | 1.85 | 1.85 |
| Cas9 | per the calculation | **0.16** |
| Mix total volume (after step 47) | 10.5 | 10.5 |

**Pre-load Cas9 with sgRNAs**

47| Add the Cas9 buffer and Cas9 to sgRNA PCR tube. Mix well, incubate at 37°C for 15 min.

**CRITICAL:** Keep PCR machine set to 37°C, pausing briefly to remove tube to add cDNA, ensuring the tube remains at 37°C.

**Cas9 Reaction**

48| Add 8 µL of purified cDNA (from step 43) to the DASH reaction (final volume is 18.5 μL). Quickly mix and return to 37°C. Incubate at 37°C for 2 h.

**Proteinase K Treatment**

49| Add 1 µL of Proteinase K to the DASH reaction. Quickly mix and return to 37°C. Incubate at 37°C for 15 min.

50| Remove the reaction from the PCR machine. Add 0.2 µL of 100 mM PMSF. Mix well.

### PCR cleanup (45 min)

51| Transfer the 19.7 µL reaction into a new Eppendorf tube, add 5.3 μL of nuclease-free water and 1.4X RNAclean XP beads (35 µL), mix by pipetting 15X and incubate the tube at room temperature for 15 min.

52| Place the tube on a magnetic rack for about 5 min until solution is clear and discard the supernatant.

53| Wash the beads with 200 µL of freshly prepared 80% (vol/vol) ethanol without removing the tube from the magnetic rack. Incubate the tube for 30 s and discard the supernatant. Repeat the wash once more and discard all the remaining ethanol droplets. Leave the tube open to air-dry for 5 min.

54| Resuspend the dried beads in 12 µL of nuclease-free water. Incubate the tube for 2 min at room temperature and then place the tube on the magnetic rack. **SAVE LIQUID** After the solution appears clear, transfer the supernatant to a new Eppendorf tube.

**POSSIBLE PAUSE:** Store the amplified libraries at −80 °C for up to 1 month or proceed to the next step.

_________________________________________________________________________________

**DAY 4**

### PCR enrichment test to determine the final number of PCR cycles (45 min)

55| Prepare the following PCR mix:

| **Reagent** | **Volume (****µL) per reaction** |
| --- | --- |
| Nuclease-free water | 3.5 |
| P5_Enr primer (12.5 µM) | 2 |
| P7_BC1_Enr primer (12.5 µM) | 2 |
| cDNA | 5 |
| KAPA HIFI HotStart ReadyMix, 2× | 12.5 |
| Mix total volume | 25 |

**CRITICAL:** To increase multiplexing, it is possible to use P7 enrichment primers that carry different indexes. An additional primer is included in Table 1 of PMID: 29215635.

56| Prepare 8-µL aliquots of PCR mix in three PCR tubes.

57| Amplify the cDNA using 5/7/9 cycles with the following PCR program. Also, set up a NO Template Control (NTC) in 9 cycles.

| **Step** | **Temperature (°C)** | **Time** | **Number of cycles** |
| --- | --- | --- | --- |
| Initial denaturation | 95 | 3 min | 1 |
| Denaturation | 95 | 30 s |  |
| Annealing | 55 | 30 s | 5/7/9 |
| Extension | 72 | 30 s |  |
| Final extension | 72 | 10 min | 1 |
| Hold | 4 | ∞ |  |

**POSSIBLE PAUSE:** Store the amplified libraries at −80°C for up to 1 month or proceed to the next step.

### PCR cleanup (1.5 h)

58| Add 17.0 μL of nuclease-free water to a final volume of 25 µL. Transfer into a new tube, add 1.4X RNAclean XP beads (35 µL), mix by pipetting 15X and incubate the tube at room temperature for 15 min.

59| Place the tube on a magnetic rack for about 5 min until solution is clear and discard the supernatant.

60| Wash the beads with 200 µL of freshly prepared 80% (vol/vol) ethanol without removing the tube from the magnetic rack. Incubate the tube for 30 s and discard the supernatant. Repeat the wash once more and discard all the remaining ethanol droplets. Leave the tube open to air-dry for 5 min.

61| Resuspend the dried beads in 12 µL of 1X Low TE. Incubate the tube for 2 min at room temperature and then place the tube on the magnetic rack. **SAVE LIQUID** After the solution appears clear, transfer the supernatant containing the eluted PCR fragments to a new Eppendorf tube.

**POSSIBLE PAUSE:** Store the amplified libraries at −80 °C for up to 1 month or proceed to the next step.

**DNA concentration and size analysis by Qubit and Agilent TapeStation (or can use BioAnalyzer)**

62| Analyze 1 µL of each PCR product using the Qubit dsDNA HS Assay Kit (Invitrogen, cat. no. Q32854), following the manufacturer’s instructions. Estimate the PCR product concentration (ng/µL), which should increase with more cycles and none with the NTC. The typical concentration range is 2–70 ng/µL.

63| Dilute 1 µL of each PCR product and anlayze using Agilent TapeStation and High-Sensitivity D1000 ScreenTape (Agilent Technologies, cat. no. 5067-5584). Estimate the average size of the PCR products. A representative profile is shown in Figure 5 of PMID: 29215635.

**CRITICAL:** The minimal library fragment size is 185 bp (insert size is ~50 bp). Smaller fragments represent incomplete amplification products, primer dimers or fragments that will yield very short reads. Therefore, if the libraries contain products smaller than 185 bp, the sample should be cleaned up once more with 1× RNAClean XP beads, as described in Steps 58–61, and the Qubit and TapeStation analyses should be redone.

**POSSIBLE PAUSE:** Store the amplified libraries at −80 °C for up to 5 years or proceed to the next step.

**SEQUENCING**

64| Depending on the cDNA profile and concentration requirements for the type of RNA sequencing performed (NextSeq, HiSeq, NovaSeq, etc.), one of the 5/7/9 cycles (from Step 61) may be sufficient to sequence. Fewer PCR cycles are better, to reduce amplification bias.

**CRITICAL:** If additional PCR amplication is required, complete this additional PCR step.

### PCR amplification of the libraries (1 h)

65| Choose one optimal PCR cycle number based on the analysis described in Steps 63.

66| Set up the following PCR mix:

| **Reagent** | **Volume (µL) per reaction** |
| --- | --- |
| Nuclease-free water | 1.5 |
| P5_Enr primer (12.5 µM) | 2 |
| P7_BC1_Enr primer (12.5 µM) | 2 |
| cDNA | 7 |
| KAPA HIFI HotStart ReadyMix, 2× | 12.5 |
| Mix total volume | 25 |

67| Mix well and amplify the cDNA using the chosen number of cycles (*x*) with the following PCR program:

| **Step** | **Temperature (°C)** | **Time** | **Number of cycles** |
| --- | --- | --- | --- |
| Initial denaturation | 95 | 3 min | 1 |
| Denaturation | 95 | 30 s |  |
| Annealing | 55 | 30 s | X |
| Extension | 72 | 30 s |  |
| Final extension | 72 | 10 min | 1 |
| Hold | 4 | ∞ |  |

**POSSIBLE PAUSE:** Store the sample at −80 °C for up to 5 years or proceed to the next step.

### PCR cleanup (1 h)

68| Transfer into a new tube, add 1.4X RNAclean XP beads (35 µL), mix by pipetting 15X and incubate the tube at room temperature for 15 min.

69| Place the tube on a magnetic rack for about 5 min until solution is clear and discard the supernatant.

70| Wash the beads with 200 µL of freshly prepared 80% (vol/vol) ethanol without removing the tube from the magnetic rack. Incubate the tube for 30 s and discard the supernatant. Repeat the wash once more and discard all the remaining ethanol droplets. Leave the tube open to air-dry for 5 min.

71| Resuspend the dried beads in 12 µL of 1X Low TE. Incubate the tube for 2 min at room temperature and then place the tube on the magnetic rack. **SAVE LIQUID** After the solution appears clear, transfer the supernatant containing the eluted PCR fragments to a new Eppendorf tube.

72| Analyze the amplified cDNA using Agilent TapeStation and Qubit as described in Steps 62–63.

**CRITICAL:** If the libraries contain products smaller than 185 bp, the sample should be cleaned up once more with 1X RNAClean XP beads.

The amplified libraries can be stored at −80 °C for up to 5 years.
