## Supplemental Protocol File 1 for "RNA-RNA Interactome Approaches Provide *in vivo* Evidence for a Critical Role of the Hfq Rim Face in sRNA-mRNA Pairing"

**Protocol for RIL-seq (with DASH)**

For RIL-seq, RNAs are crosslinked to an RNA binding protein of interest, protein is immunoprecipitated (using native antibodies or antibodies against tag on protein), RNAs associated with the protein are trimmed and then ends that are in proximity are ligated. In the protocol described here, rRNA is depleted using DASH.

This protocol was adapted and optimized based on the following previously published methods:

- The RIL-seq protocol described in PMID: 29215635.
- rRNA depletion was performed using DASH (Depletion of Abundant Sequences by Hybridization), modified from PMID: 32345633.
- Barcode sequences are described in PMID: 25730492 (where additional sequences can be found).

_________________________________________________________________________________

**DAY 1**

**Cell growth, cross-linking and freezing of bacterial pellets (24 h)**

**CRITICAL:** In the following steps, the growth conditions (growth phase, medium and temperature, as well as different stress conditions) can be varied. Here, we describe the protocol for an experiment with

*E. coli* grown to log phase in LB medium at 37 °C with shaking.

1| Grow 2 mL of overnight cultures of strains, starting from single colonies under desired conditions. The next day, dilute the overnight cultures 100-fold in fresh medium and grow to desired OD_600nm_. The culture volume to be grown is calculated according to the equation ‘culture volume = 40/OD_600nm_ value’

(e.g., 80 mL for cells at OD_600nm_ value of 0.5).

2| Transfer the culture to 50-mL tubes and centrifuge the tubes at 4,500*g* for 10–15 min at 4 °C. Note that you may need more than one 50-mL tube per sample. Combine the tubes of the same sample in the next washing step (Step 3).

3| Discard the supernatants, resuspend the pelleted cells in 20 mL of ice-cold 1X PBS and centrifuge at 4,500*g* for 10 min at 4 °C. Repeat the PBS wash and resuspend the pelleted cells in 10 mL of ice-cold 1X PBS.

**CRITICAL:** From this step and until the cells are being frozen in liquid nitrogen (Step 9), samples must be kept on ice at all times.

4| Spread the 10-mL suspension of one sample in an empty Petri dish (plastic) and immediately place the dish on a metal block cooled to −20 °C.

5| Expose the cells to 80,000 µJ/cm^2^ of 254-nm UV irradiation, at a distance of 3 cm from the bulbs, using

a Stratalinker 1800 UV cross-linker.

6| Transfer the irradiated cells to a cold 50-mL tube and immediately place it on ice.

**CRITICAL:** Keep the treated and pretreated samples on ice while treating the other samples.

7| Centrifuge the cell suspensions at 4,500*g* for 10 min at 4 °C. Resuspend the cells in 1 mL of ice-cold PBS and transfer the suspensions to precooled 2-mL Eppendorf tubes.

8| Centrifuge the cell suspensions at 17,000*g* for 3 min at 4 °C and discard the supernatants.

9| Freeze the pellets using liquid nitrogen and store them at −80 °C.

**POSSIBLE PAUSE:** Frozen pellets can be kept at –80 °C for at least 1 week.

__________________________________________________________________________________

**DAY 2**

**Lysate preparation (1-2 h)**

**CRITICAL:** Work on ice and use cold (4 °C) wash buffer during the lysate preparation.

10| Prepare RNase inhibitor-free wash buffer.

| **Reagent** | **Volume (µL) per reaction** | **Final concentration** | **Per 14 samples (µL)** |
| --- | --- | --- | --- |
| Salts solution for wash buffer * | 4,920 |  | 68,880 |
| IGEPAL 100% (wt/vol)  (Sigma-Aldrich, cat. no. I8896) | 5 | 0.1% | 70 |
| Imidazole, 1 M | 50 | 10 mM | 700 |
| Protease inhibitor cocktail | 25 |  | 350 |
| Buffer total volume | 5,000 |  | 70,000 |

* Salts solution for RNAse inhibitor-free wash buffer = 250 µL 1M sodium phosphate buffer pH 8.0 mix** + 300 µL 5M NaCl + 4.37 mL DEPC H_2_O

** Prepare 1M Sodium Phosphate buffer pH 8.0 Mix (mix 6.8 mL of sterile 1M Na_2_H_2_PO_4_ and 93.2 mL of sterile 1M Na_2_HPO_4_ in a sterile bottle. This buffer can be stored at room temperature for a year or more)

**CRITICAL:** It may take a few min for the IGEPAL to dissolve; make sure that it is fully dissolved (by vortexing) before starting to work with the buffer.

11| Withdraw 0.5 mL of the RNase inhibitor-free wash buffer per sample and keep it on ice for Step 26. To the remaining buffer (4.5 mL per sample), add 11.25 µL of recombinant RNase inhibitor (40 U/µL) for a final concentration of 0.1 U/µL. This is the wash buffer. Keep the buffers on ice.

12| For each sample, prepare a new 2-mL Eppendorf tube filled with 0.1-mm diameter glass beads up to the
400-µL mark. Place the tubes on ice.

13| Thaw the frozen pelleted cells from Step 9 on ice, and resuspend the cells with 800 µL of cold wash buffer by pipetting up and down. Transfer the resuspended cells to the 2-mL tubes prepared in Step 12.

14| Vortex 30 sec, 10X. Rest 30 sec on ice between each vortexing.

15| Centrifuge the tubes at 14,000*rpm* for 3 min at 4 °C. Transfer the supernatants (600 µL) to new Eppendorf tubes. This is the bacterial lysate.

**CRITICAL:** You can leave a small fraction of the supernatant in the grinding tube, to avoid transferring the glass beads.

16| Add 400 µL of cold wash buffer to the grinding tubes containing the glass beads and vortex 1x for

30 sec.

17| Centrifuge the tubes at 14,000*rpm* for 3 min at 4 °C and transfer the supernatants (400 µL) to the tubes already containing the lysate from Step 15.

18| Centrifuge the collected lysate (from Step 15 and 17) at 14,000*rpm* for 15 min at 4 °C. Transfer the lysates to new Eppendorf tubes. The total volume of the lysate should be ~800–900 µL.

**CRITICAL:** The lysate should be kept on ice and used within a short time on the same day in the following IP procedure.

19| Transfer 75 µL of the lysate of each sample from Step 18 to a new tube and keep the tube on ice. Add 225 µL wash buffer (total 300 µL), followed by 900 µL TriReagent (Sigma-Aldrich, cat. no. T9424). Pipette 10X and store at -80°C for total RNA extraction. Extract the total RNA following the standard TriReagent protocol, then construct cDNA libraries for RNA-seq using the Protocol for RNA-seq (with DASH); Supplementary Protocol File 2).

Also, keep another 24 µL of lysate for Western blot, adding preferred loading buffer.

##### Lysate pre-clearing (1.5 h)

20| Mix the protein A/G magnetic bead solution by inversion or gentle vortexing. For each sample, aliquot
20 µL of beads in a new tube. Add 200 µL of cold wash buffer, mix by tapping the tubes and place the tubes on a magnetic rack. After the solution becomes clear, discard the buffer while the tubes are still on the rack. Remove the tubes from the magnetic rack. Add 750 µL of cell lysate from Step 18 to the washed magnetic beads and rotate the mixture for 60 min at 4 °C. In the meantime, proceed with Step 21.

**CRITICAL:** Ensure that all the beads adhere to the magnet and that the solution is clear before removing the buffer.

##### Co-immunoprecipitation using anti-Flag antibody (or native antibodies) (3 h)

**CRITICAL:** This protocol assumes the use of Flag-tagged binding proteins. If the study is not being carried out with a Flag-tagged RNA binding protein, the antibody volume should be adjusted according to the antiserum used, and optimization may be necessary.

**CRITICAL:** If possible, Steps 21–25 should be performed in a cold room. Moving the samples from 4 °C to room temperature may reduce the efficiency of the protocol.

**CRITICAL:** From this step on, it is highly recommended to use LoBind or Maxymum Recovery tubes in order to decrease RNA loss.

21| For each sample, aliquot 20 µL of well-mixed protein A/G magnetic beads in a new tube. Add 200 µL of cold wash buffer, mix by tapping the tubes and place the tubes on a magnetic rack. After the solution becomes clear, discard the buffer while the tubes are still on the rack, without touching the beads. Add 200 µL of cold wash buffer and 3 µL of anti-Flag M2 antibody to the washed beads. Rotate the mixture for 30 min at 4 °C to bind the anti-Flag M2 antibody to the protein A/G magnetic beads.

22| Spin down the tubes from Step 21 and place them on a magnetic rack. After the solution becomes clear, discard the supernatant containing unbound antibody while the tubes are on the magnetic rack. Remove the tubes from the rack, add 200 µL of cold wash buffer, mix by tapping the tubes and place the tubes on the magnetic rack. After the solution becomes clear, discard the buffer while the tubes are on the magnetic rack. Remove the tubes from the magnetic rack and place them on ice.

23| Spin down the tubes from Step 20, which containing protein A/G beads and lysate, and place them on a magnetic rack for 5 min. After the solution becomes clear, transfer the cleared lysate to the tubes prepared in Step 22. Rotate the tubes for 90 min at 4 °C to bind the Flag-tagged protein to the anti-Flag antibodies on the magnetic beads.

24| Spin down the tubes from Step 23 at 1,500*g* for 10 s at room temperature and place them on a magnetic rack for 5 min. After the solution becomes clear, discard the lysate and remove the tubes from the magnetic rack.

25| Wash the beads: add 200 µL of cold wash buffer and rotate the tubes for 10 min at 4 °C. Spin down the tubes and place them on a magnetic rack for 1 min. After the solution becomes clear, discard the buffer. Repeat the washes four additional times. In the meantime, prepare the buffers described in Steps 26 and 27.

##### Trimming RNA ends (40 min)

26| Prewarm the RNase inhibitor-free wash buffer that was withdrawn in Step 11 (500 µL per sample) to room temperature. Dilute the RNase A/T1 mixture 1/20 in water (1 µL mixture and 19 µL of DEPC H_2_O). Mix 1 µL of diluted RNase A/T1 with 499 µL of prewarmed RNase inhibitor-free wash buffer to prepare 500 µL of RNase digestion buffer per sample. This volume includes 20 µL extra, in order to have sufficient amount for all the samples.

**CRITICAL:** When pipetting the RNase A/T1 mixture for the preparation of the 1/20 dilution, make sure not to have an additional amount of enzyme outside the tip, as this would increase the RNase concentration.

27| Prepare 650 µL of SUPERase IN Wash per sample by adding 3.25 µL of SUPERase IN RNase inhibitor to 647 µL of wash buffer, resulting in a final concentration of 0.1 U/µL. Keep the buffer on ice. SUPERase IN RNase inhibitor also inhibits RNase T1, which is not inhibited by the recombinant RNase inhibitor.

28| To trim the exposed parts of the RNAs, add 480 µL of RNase digestion buffer prepared in Step 26 to the tubes from Step 25. Incubate the tubes at 22 °C with gentle agitation for 7 min, 300 rpm.

**CRITICAL:** If the magnetic beads collect at the bottom of the tubes during the incubation time, you can tap the tubes a few times or vortex very gently for 2–3 s to obtain a homogenous suspension.

**CRITICAL:** Note that extended incubation of the samples with the RNase A/T1 mixture may result in overdigestion of the RNA.

29| Place the samples briefly on ice for 30 sec to slow the reaction, spin down the tubes at 1,500*g* for 10 s at room temperature and then place them on a magnetic rack. After the solution becomes clear, remove the RNase digestion buffer. Add 200 µL of cold SUPERase IN Wash prepared in Step 27 to each tube, rotate the tubes for 5 min at 4 °C and then spin down the tubes and place them on the magnetic rack. After the solution becomes clear, discard the buffer. Repeat the wash an additional 2X. In the last wash, do not discard the buffer and store the tubes on ice until Step 31.

**CRITICAL:** The washes in Step 29 should be performed in a cold room.

##### 5′OH end phosphorylation and 2′P/3′P end dephosphorylation (2.5 h)

**CRITICAL:** In Steps 30–32, the 5′OH and 2′P/3′P RNA ends that were formed by the nuclease treatment (Steps 26–29) are modified by T4 polynucleotide kinase (PNK), which is capable of both phosphorylating the
5′ OH end and dephosphorylating the 2′P/3′P ends of the RNA. Thus, PNK generates fragments with 5′P and 3′OH ends that can be subsequently ligated (Step 33).

30| Prepare the following PNK reaction mix (80 µL per sample; *x* samples+1 µL extra):

| **Reagent** | **Volume (µL) per reaction** | **Final concentration** | **Per 14 samples (µL)** |
| --- | --- | --- | --- |
| Nuclease-free water | 65.2 |  | 912.8 |
| PNK buffer, 10× | 8 | 1X | 112 |
| ATP (100 mM)a | 0.8 | 1 mM | 11.2 |
| Recombinant RNase inhibitor (40 U/µL) | 2 | 1 U/µL | 28 |
| T4 polynucleotide kinase (10,000 U/ml) | 4 | 0.5 U/µL | 56 |
| Mix total volume | 80 |  | 1,120 |

31| Place the tubes from Step 29 on a magnetic rack and, after the solution becomes clear, discard the wash buffer. Remove the tubes from the magnetic rack and add 80 µL of PNK reaction mix to each tube. Incubate the tubes for 2 h at 22 °C with a gentle agitation, 300 rpm.

**CRITICAL:** If the magnetic beads precipitate during the incubation time, vortex them very gently for 2–3 s once every 30 min.

32| Place the tubes on a magnetic rack and, after the solution becomes clear, remove the PNK reaction mixture and add 200 µL of cold wash buffer to each tube. Rotate the tubes for 5 min at 4 °C. Place the tubes on the magnetic rack and, after the solution becomes clear, discard the buffer. Repeat the washes with cold wash buffer two additional times. To keep the beads from drying out, retain the wash buffer from the last wash until the ligation mixture is ready in the next step and remove it before the addition of the ligation mixture.

##### Ligation of neighboring RNAs (overnight)

33| Set up the following ligation mix (80 µL per sample; *x* samples + 2 µL) at room temperature:

| **Reagent** | **Volume (µL) per reaction** | **Final concentration** | **Per 15 samples (µL)** |
| --- | --- | --- | --- |
| 10× T4 RNA ligase buffer | 8 |  | 120 |
| DMSO (100% (vol/vol)) | 7.2 |  | 108 |
| ATP (100 mM) | 0.8 | 1 mM | 12 |
| PEG 8000 (50% (wt/vol)) | 32 |  | 480 |
| Recombinant RNase inhibitor (40 U/µL) | 1.2 | 0.6 U/µL | 18 |
| T4 RNA ligase 1 (30,000 U/mL) | 7.2 | 2.7 U/µL | 108 |
| Nuclease-free water | 23.6 |  | 354 |
| Mix total volume | 80 |  | 1,200 |

**CRITICAL:** Set up the mix at room temperature to prevent DMSO precipitation. Pipette the PEG very slowly for accurate aspiration, as it is very viscous. Mix well by tapping the tube, as the mixture is very viscous, and spin down.

34| Add 80 µL of ligation mixture to each tube. Mix well by tapping the tube, and if necessary spin down.

35| Incubate the tubes at 22 °C overnight, without agitation, covered with foil.

__________________________________________________________________________________

**DAY 3**

##### Protein digestion (2.5 h)

36| Prepare fresh wash buffer (the amounts per single sample are detailed):

| **Reagent** | **Volume (µL) per reaction** | **Final concentration** | **Per 14 samples (µL)** |
| --- | --- | --- | --- |
| Salts solution for wash buffer* | 638 |  | 8,932 |
| IGEPAL 100% (wt/vol) | 0.65 | 0.1% | 9.1 |
| Imidazole 1 M | 6.5 | 10 mM | 91 |
| Recombinant RNase inhibitor (40 U/µL) | 1.625 | 0.1 U/µL | 22.75 |
| Protease inhibitor cocktail | 3.25 |  | 45.5 |
| Buffer total volume | 650 |  | 9,100 |

* Prepare 10 mL Salts solution for wash buffer = 0.5 mL 1M Sodium Phosphate buffer pH 8.0 mix** + 0.6 mL 5M NaCl + 8.72 mL DEPC H_2_O

** Prepare 1M sodium phosphate buffer pH 8.0 Mix (mix 6.8 mL of sterile 1M Na_2_H_2_PO_4_ and 93.2 mL of sterile 1M Na_2_HPO_4_ in a sterile bottle. This buffer can be stored at room temperature for a year or more)

**CRITICAL:** Keep the buffer on ice.

37| Place the tubes from Step 35 in the magnetic rack and allow the solution to become clear. This may take a few minutes because of the viscosity of the solution.

38| After the solution becomes clear (during the next 10 min), make proteinase K buffer and prepare fresh proteinase K reaction mix (300 µL per sample; *x* samples +1 µL extra) by mixing the following components at room temperature:

| **Reagent** | **Volume (µL) per reaction** | **Final concentration** | **Per 14 samples (µL)** |
| --- | --- | --- | --- |
| Nuclease-free water | 240 |  | 3,360 |
| Tris-HCl, pH 7.8 (1 M)  [400 mL Tris HCl, pH 8.0 and 400 mL Tris HCl, pH 7.5] | 15 | 50 mM | 210 |
| NaCl (5 M) | 3 | 50 mM | 42 |
| IGEPAL (100% (wt/vol)) | 0.3 | 0.1% | 4.2 |
| Imidazole (1 M) | 3 | 10 mM | 42 |
| SDS (10% (wt/vol)) | 30 | 1% | 420 |
| EDTA pH 8.0 (0.5 M) | 3 | 5 mM | 42 |
| β-mercaptoethanol (14.3 M) | 0.1 | 5 mM | 1.4 |
| Recombinant RNase inhibitor (40 U/µL) | 1.3 | 0.1 U/µL | 18.2 |
| Proteinase K (20 mg/ml) | 5 |  | 70 |
| Mix total volume | 300 |  | 4,200 |

! β-mercaptoethanol is toxic. The stock should be handled in a fume hood, while working with appropriate protective equipment.

39| Discard the ligation mix (leave a small amount in for first wash – it prevents loss of beads) and add 200 µL of cold wash buffer prepared in Step 36 to each tube. Rotate the tubes for 5 min at 4 °C. Spin down the tubes and place them on the magnetic rack. After the solution becomes clear, discard the buffer. Repeat the washes with cold wash buffer two additional times. (COLD ROOM)

40| Add 300 µL of proteinase K reaction mix to each tube. Mix well and incubate the samples for 2 h at 55°C with gentle agitation 300 rpm.

**CRITICAL:** If the magnetic beads precipitate during the incubation, vortex them very gently for 2–3 sec once every 30 min.

##### RNA extraction according to the standard TriReagent LS protocol (14 h)

**CRITICAL:** From this step and until the end of the experimental procedure, use filter tips only.

41| Add 0.9 mL of TriReagent LS (Sigma-Aldrich, cat. no. T3934) (prewarmed to room temperature) to the tubes containing both the beads and the proteinase K buffer. Resuspend thoroughly by pipetting to homogenization, and incubate the tubes for 5 min at room temperature.

! TriReagent LS is toxic and should be handled in a fume hood, while working with appropriate protective equipment.

42| Add 200 µL of chloroform, mix by inversion of the tubes for 15 s and then incubate them for 10 min at room temperature.

! Chloroform is toxic and should be handled in a fume hood, while working with appropriate protective equipment.

43| Centrifuge the tubes at 17,000*g* for 10 min at 4 °C and transfer the upper phase (~700 µL) to a new Eppendorf tube.

! The lower phase contains phenol and should be disposed of as toxic waste.

**CRITICAL:** Be careful not to transfer the lower phase to the new tube.

44| Add 500 µL of isopropanol, mix thoroughly by inversion of the tubes and incubate them for 10 min at room temperature.

45| Spin down the tubes, add 1.5 µL of GlycoBlue and mix well by inverting tubes. Incubate the tubes overnight at −20 °C.

**CRITICAL:** As the RNA amount at this step is very small, the addition of GlycoBlue is critical to precipitating the RNA.

**POSSIBLE PAUSE:** Samples can be stored at −20 °C for up to 1 month before proceeding to the next step.

__________________________________________________________________________________

**DAY 4**

46| Centrifuge the tubes at 14,000*rpm* for 15 min at 4 °C and discard the supernatant.

47| Wash the pellet by the addition of 1 mL of freshly made 75% (vol/vol) ethanol, followed by centrifugation at 12,000*rpm* for 5 min at 4 °C and discarding of the supernatant. Repeat this washing step once. After discarding the supernatant from the second wash, spin down the tubes again and discard the remaining supernatant. Leave the tubes open for 3 min to dry the pellet.

**CRITICAL:** When performing the washes, be careful not to lose the RNA pellet.

48| Resuspend the pellet in 20 µL of nuclease-free water by tapping the tube. If the resuspension is difficult, warm the tube at 37 °C for a few min and flick the tube. After the resuspension, place the RNA immediately on ice.

**CRITICAL:** It is important not to resuspend the pellet by pipetting, as the amount of the RNA is very small and the pellet might stick to the tip.

49| Analyze RNA samples (not diluted) from Step 48 using a Agilent TapeStation (or use BioAnalyzer) to assess RNA quantity and size distribution. Typically, the concentrations obtained for the samples from Step 48 range between 500 and 1,500 ng/µL.

50| Transfer 15 µL of each RNA solution from Step 48 to PCR tubes and add 1 μL of recombinant RNase inhibitor (40 U/µL).

**POSSIBLE PAUSE:** Continue with RNA-seq library construction or store the samples at −80 °C for up to 1 month.

**RNA-seq library construction and rRNA removal via DASH**

__________________________________________________________________________________

**DAY 5**

### Fragmentation and DNase–FastAP combined treatment (45 min)

**CRITICAL:** RIL-seq libraries are constructed using the RNAtag-Seq protocol with a few modifications to allow capture of short RNA fragments. The library construction process includes a step of random RNA fragmentation. This step is required to obtain full coverage of long RNAs.

61| Add 11 µL of the eluted sample from Step 60 to a new PCR tube and add 1 µL of 50 µM AR2 primer (sequence: 5’-TACACGACGCTCTTCCGAT-3’; also refer to Table 1 of PMID: 29215635). Mix well.

62| Heat the mixture to 70 °C for 2 min and immediately place it on ice.

63| Set up the following reverse transcription mix on ice:

| **Reagent** | **Volume (µL) per reaction** | **Final concentration (25 µL)** |
| --- | --- | --- |
| dNTP mix (10 mM) | 1.25 | 0.5 mM |
| 10× RT buffer | 2.5 |  |
| MgCl2 (25 mM) | 5 | 5 mM |
| DTT (0.1 M) | 2.5 | 0.01 M |
| RNaseOUT (40 U/µL) | 0.5 | 0.8 U/µL |
| SuperScript III RT (200 U/µL) | 1.25 | 10 U/µL |
| Mix total volume | 13 |  |

64| Add 13 µL of reverse transcription mix to the mixture from Step 62. Mix well and incubate the mixture at 50 °C for 55 min.

**POSSIBLE PAUSE:** Store the sample at −80°C for up to 1 month or proceed to the next step.

### RNA degradation (20 min)

65| Add 2.5 µL of 1N NaOH (2 µL of 5N NaOH and 8 µL DEPC H_2_O) to the tube from Step 64 and incubate it at 70 °C for 12 min.

66| Add 5 µL of freshly diluted 0.5 M acetic acid (1/33 from original stock (16.5 M from Sigma) – 30 µL of stock + 970 µL of H_2_O) and mix well.

### cDNA cleanup (45 min)

67| Add 7.5 µL of nuclease-free water for a final volume of 40 µL and transfer it to a new Eppendorf tube.

68| Add 1.5X (60 µL) isopropanol and 2.5X (100 µL) RNAclean XP beads (Beckman Coulter, cat. no. A63987), mix by pipetting 15X and incubate the tube at room temperature for 15 min.

69| Place the tube on a magnetic rack for about 5 min until the solution is clear and discard the supernatant.

70| Wash the beads with 200 µL of freshly prepared 80% (vol/vol) ethanol without removing the tube from the magnetic rack. Incubate the tube for 30 s and discard the supernatant. Repeat the wash once more, and discard all the remaining ethanol droplets.

71| Leave the tube open on the magnetic rack and allow the beads to air-dry at room temperature for 10 min.

72| Remove the tube from the magnetic rack, resuspend the beads in 5 µL of nuclease-free water and incubate the tube for 2 min at room temperature.

**CRITICAL:** Keep the cDNA in the tube with the beads for the following ligation step.

### Ligation of a second adaptor at the cDNA 3’ end (ssDNA/ssDNA) (overnight)

73| Transfer DNA with beads to a new PCR tube and add 2 µL of 40 µM 3Tr3 adaptor (sequence: P-AGATCGGAAGAGCACACGTCTG-ddC; also refer to Table 1 of PMID: 29215635) to the cDNA+beads solution, incubate the tube at 75°C for 3 min and place it on ice.

74| Set up the following ligation mix at room temperature:

| **Reagent** | **Volume (µL) per reaction** | **Final concentration (20 µL)** |
| --- | --- | --- |
| T4 RNA ligase buffer, 10× | 2 |  |
| DMSO (100% (vol/vol)) | 0.8 |  |
| ATP (100 mM) | 0.2 | 1 mM |
| PEG 8000 (50% (wt/vol)) | 8.5 |  |
| T4 RNA ligase 1 (30,000 U/mL) | 1.5 | 2.25 U/mL |
| Mix total volume | 13 |  |

**CRITICAL:** Set up the mix at room temperature to prevent DMSO precipitation. Pipette very slowly for accurate aspiration of PEG, as it is very viscous. Mix well by tapping the tube, as the mixture is very viscous, and spin down.

75| Add 13 µL of ligation mix to the tube from Step 73. Mix well by tapping the tubes, as the solution is very viscous. Spin down and incubate the tube overnight at 22°C.

_________________________________________________________________________________

**DAY 7**

### Cleanup of cDNA (45 min)

76| Increase the volume to 40 µL by adding 20 µL of nuclease-free water and transfer to a new Eppendorf tube. Add 1.5X (60 µL) isopropanol and 2.5X (100 µL) RNAClean XP beads, mix by pipetting 15X and incubate the tube at room temperature for 15 min.

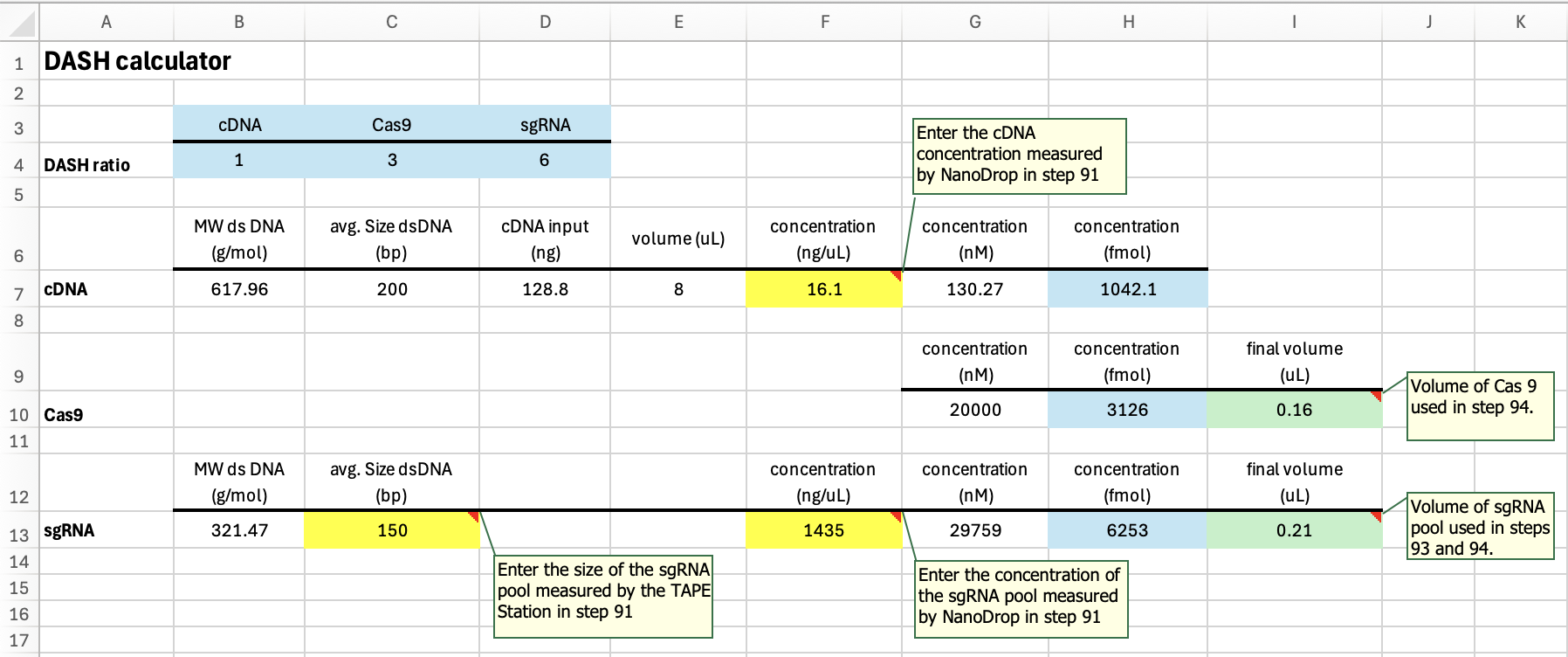

93| Add calculated sgRNA (ex. 0.21 μL) and nuclease-free water (amount per the table below) to a new PCR tube, incubate the tube at 94°C for 3 min, 4°C for 5 min.

94| Prepare the DASH reaction:

| **Reagent** | **Volume (****µL ) per reaction** | **Example reaction (µL)** |
| --- | --- | --- |
| sgRNA pool | per the calculation | **0.21** |
| Nuclease-free water | up to 10.5 | 8.28 |
| Cas9 buffer (3.1 NEB buffer) 10x | 1.85 | 1.85 |
| Cas9 | per the calculation | **0.16** |
| Mix total volume (after step 95) | 10.5 | 10.5 |

**Pre-load Cas9 with sgRNAs**

95| Add the Cas9 buffer and Cas9 to sgRNA PCR tube. Mix well, incubate at 37°C for 15 min.

**CRITICAL:** Keep PCR machine set to 37°C, pausing briefly to remove tube to add cDNA, ensuring the tube remains at 37°C.

**Cas9 Reaction**

96| Add 8 µL of purified cDNA (from step 91) to the DASH reaction (final volume is 18.5 μL). Quickly mix and return to 37°C. Incubate at 37°C for 2 h.

**Proteinase K Treatment**

97| Add 1 µL of Proteinase K to the DASH reaction. Quickly mix and return to 37°C. Incubate at 37°C for 15 min.

98| Remove the reaction from the PCR machine. Add 0.2 µL of 100 mM PMSF. Mix well.

### PCR cleanup (45 min)

99| Transfer the 19.7 µL reaction into a new Eppendorf tube, add 5.3 μL of nuclease-free water and 1.4X RNAclean XP beads (35 µL), mix by pipetting 15X and incubate the tube at room temperature for 15 min.

100| Place the tube on a magnetic rack for about 5 min until solution is clear and discard the supernatant.

101| Wash the beads with 200 µL of freshly prepared 80% (vol/vol) ethanol without removing the tube from the magnetic rack. Incubate the tube for 30 s and discard the supernatant. Repeat the wash once more and discard all the remaining ethanol droplets. Leave the tube open to air-dry for 5 min.

102| Resuspend the dried beads in 12 µL of nuclease-free water. Incubate the tube for 2 min at room temperature and then place the tube on the magnetic rack. **SAVE LIQUID** After the solution appears clear, transfer the supernatant to a new Eppendorf tube.

**POSSIBLE PAUSE:** Store the amplified libraries at −80 °C for up to 1 month or proceed to the next step.

_________________________________________________________________________________

**DAY 8**

### PCR enrichment test to determine the final number of PCR cycles (45 min)

103| Prepare the following PCR mix:
