## Supplemental Protocol File 3 for "RNA-RNA Interactome Approaches Provide *in vivo* Evidence for a Critical Role of the Hfq Rim Face in sRNA-mRNA Pairing"

**Protocol for Hi-GRIL-seq and iRIL-seq (with DASH)**

For Hi-GRIL-seq and iRIL-seq, T4 RNA ligase is briefly expressed in cells (sufficiently long to ligate 3’ and 5’ RNA ends that are in proximity but briefly enough to prevent cell death). For Hi-GRIL-seq, total RNA is isolated. For iRIL-seq, RNA that co-associates with a specific RNA binding protein is isolated. In the protocol described here, rRNA is depleted using DASH.

This protocol was adapted and optimized based on the following previously published methods:

- In vivo RNA ligation steps were modified from protocols described in PMID: 28976035 and PMID: 38062076.
- cDNA library construction was adapted from the RIL-seq protocol described in PMID: 29215635.
- rRNA depletion was performed using DASH (Depletion of Abundant Sequences by Hybridization), modified from PMID: 32345633.
- Barcode sequences are described in PMID: 25730492 (where additional sequences can be found)

_________________________________________________________________________________

**DAY 1**

### Bacterial cultures and *in vivo* ligation (3 h)

1| Pick a single colony of *E. coli* strains harboring either pBAD24 (vector control) or pBAD24-*t4ml1* (T4 RNA Ligase I expression plasmid). Inoculate into 10 mL of LB (Lennox) supplemented with 100 µg/mL ampicillin. Grow overnight at 37 °C with shaking (~250 rpm).

2| Dilute the overnight cultures into 80 mL of fresh LB to a starting OD₆₀₀ of 0.05. Incubate at 37 °C with shaking.

3| When cultures reach OD₆₀₀ ≈ 0.5, add 0.2% arabinose to induce T4 RNA Ligase I expression.

4| When OD₆₀₀ reaches 1.0, harvest 20 OD₆₀₀ units of cells by centrifugation at 4,300 rpm for 10 min at 4 °C.

**CRITICAL:** It is best to determine the optimal induction and growth time/conditions for the specific strain and growth media being used, as these may vary depending on the experimental setup.

5| Discard most of the supernatant, leaving approximately 1 mL. Resuspend the pellet thoroughly and transfer the cell suspension to a 2 mL microcentrifuge tube.

6| Centrifuge at 14,000 rpm for 2 min at 4 °C. Discard the supernatant completely and immediately snap-freeze the pellet in liquid nitrogen.

7| Store the frozen pellet at –80 °C for up to 1 week.

### Preparation of antibody–Protein A Sepharose (PAS) beads for Hfq IP using anti-Hfq antibody (0.5 h)

8| Weigh out 120 mg of Protein A Sepharose CL-4B beads (Amersham Biosciences, cat. no. 17-0780-01) and transfer into a 2 mL RNase-free microcentrifuge tube.

9| Add 1 mL Net2 Buffer (50 mM Tris-HCl, pH 7.4; 150 mM NaCl; 0.05% Triton X-100) to allow the beads to swell.

10| Add 100 µL anti-Hfq antiserum directly to the bead suspension.

11| Incubate on a nutator or rotator at 4 °C overnight to allow antibody binding.

_________________________________________________________________________________

**DAY 2**

### Preparation of antibody–Protein A Sepharose (PAS) beads for Hfq IP (continued) (1 h)

12| Transfer the antibody–PAS beads to a 15 mL Falcon tube. Centrifuge at 4,300 rpm for 10 min at 4 °C to pellet the beads. Discard the supernatant.

13| Add 10 mL Net2 Buffer, invert the tube 10X to wash the beads, then centrifuge again at 4,300 rpm for 10 min at 4 °C. Discard the supernatant.

14| Repeat the wash step 2X more to ensure removal of unbound antibody and contaminants.

15| After the final wash, discard the supernatant completely. Keep the beads on ice or at 4 °C until use.

**Preparation of cell lysate** **(0.5 h)**

16| Retrieve the frozen cell pellet from Step 7 and add 400 µL (~0.6 g) of 212–300 µm glass beads (Sigma, cat. no. G1277), 400 µL lysis buffer (20 mM Tris-HCl, pH 8.0; 150 mM KCl; 1 mM MgCl₂; 1 mM DTT), and 2 µL RNasin (RNase inhibitor) to the same tube.

17| Lyse the cells by vortexing for 30 sec, placing on ice for 30 sec, and repeating this cycle 10X.

18| Add 800 µL lysis buffer, then vortex for 30 sec to fully resuspend and solubilize the lysate.

19| Centrifuge at 14,000 rpm for 10 min at 4 °C; the supernatant (~1 mL) is the cleared cell lysate and should be kept on ice for immediate use.

Hfq co-IP RNA isolation for iRIL-seq analysis **(3 h)**

20| Transfer 950 µL of cleared cell lysate (from Step 19) to the antibody–PAS beads (from Step 15), then add 950 µL Net2 buffer and 5 µL RNasin to the mixture.

21| Incubate on a nutator at 4 °C for 2 h to allow binding of Hfq–RNA complexes to the beads.

22| Centrifuge at 4,300 rpm for 10 min at 4 °C, then carefully discard the supernatant.

23| Add 10 mL Net2 buffer, invert the tube 10X to wash the beads, then centrifuge again at 4,300 rpm for 10 min at 4 °C and discard the supernatant.

24| Repeat the wash step 2X more.

25| Add the following to the washed beads: 2.2 mL NET2 buffer, 250 µL of 3 M sodium acetate (NaOAc), 25 µL of 10% SDS, and 3 mL of Phenol:Chloroform:Isoamyl Alcohol (25:24:1, pH 8). Vortex for 15 sec to mix thoroughly.

26| Centrifuge at 4,300 rpm for 10 min at 4 °C, then carefully transfer 2.5 mL of the upper aqueous phase to a new 15 mL Falcon tube.

27| Add 5 µL GlycoBlue and 6.25 mL of 100% ethanol to the aqueous phase. Mix by inversion.

28| Precipitate the RNA by placing the tube at –80 °C overnight.

### Total RNA isolation for Hi-GRIL-seq analysis (2 h)

29| Transfer 50 µL of cell lysate (from Step 19) into a RNase-free 1.5 mL microcentrifuge tube. Add 1 mL of TriReagent (Sigma-Aldrich, cat. no. T9424) directly to the lysate. Mix thoroughly by pipetting up and down 10X to homogenize the sample.

30| Incubate at room temperature (RT) for 5 min to allow complete dissociation of nucleoprotein complexes.

31| Add 200 µL of chloroform to the mixture. Cap tightly and vortex vigorously for 15 sec to emulsify the phases. Incubate at RT for 3 min.

32| Centrifuge the sample at 14,000 rpm, 4 °C for 15 min. The mixture will separate into a lower red phenol-chloroform phase, an interphase, and a clear upper aqueous phase containing RNA.

33| Carefully transfer 600 µL of the upper aqueous phase to a new RNase-free 1.5 mL microcentrifuge tube. Avoid disturbing the interphase.

34| Add 500 µL isopropanol to the aqueous phase. Mix by vortexing for 10 sec.

35| Incubate the mixture at RT for 10 min or at –20 °C overnight to allow RNA precipitation.

36| Centrifuge at 14,000 rpm, 4 °C for 10 min. A white or translucent pellet should be visible at the bottom of the tube. Carefully discard the supernatant.

37| Wash the RNA pellet with 1 mL of 70% ethanol (pre-chilled if possible). Centrifuge at 14,000 rpm, 4 °C for 5 min. Discard the supernatant.

38| Air-dry the pellet at RT for 10 min. Avoid overdrying.

39| Resuspend the RNA pellet in 15 µL DEPC-treated water.

40| Quantify RNA concentration using a 1:10 dilution on a NanoDrop spectrophotometer.

41| Analyze the RNA samples (diluted 1/10) using RNA ScreenTape in the TapeStation to assess RNA quantity and size distribution.

**POSSIBLE PAUSE:** RNA samples can be stored at –80 °C for up to 1 month.

_________________________________________________________________________________

**DAY 3**

Hfq co-IP RNA isolation for iRIL-seq analysis (continued) **(2 h)**

42| Retrieve the tubes from Step 28 and centrifuge at 4,300 rpm for 1 h at 4 °C to pellet the RNA. Carefully discard the supernatant.

43| Wash the blue RNA pellet with 10 mL of 70% ethanol, then centrifuge at 4,300 rpm for 10 min at 4 °C. Carefully discard the supernatant.

44| Add 1 mL of 70% ethanol and transfer the pellet into a new 1.5 mL RNase-free microcentrifuge tube. Centrifuge at 14,000 rpm for 10 min at 4 °C. Carefully discard the supernatant.

45| Air-dry the pellet at RT for 15 min.

46| Resuspend the RNA pellet in 15 µL DEPC-treated water.

47| Measure the RNA concentration using a NanoDrop. (Do not dilute the sample prior to measurement, as RNA yield may be low)

48| Analyze the RNA samples using RNA ScreenTape in the TapeStation to assess RNA quantity and size distribution.

**POSSIBLE PAUSE:** RNA samples can be stored at –80 °C for up to 1 month.

RNA dilution preparation for cDNA library construction (1 h)

49| Using RNA isolated from Total RNA (Step 39) and Hfq IP (Step 46), calculate the volume required to obtain up to 400 ng of RNA based on the measured concentrations.

50| Adjust the final volume to 15 µL with DEPC-treated water in RNase-free PCR tubes. Add 1 µL of recombinant RNase inhibitor (40 U/µL) to each tube.

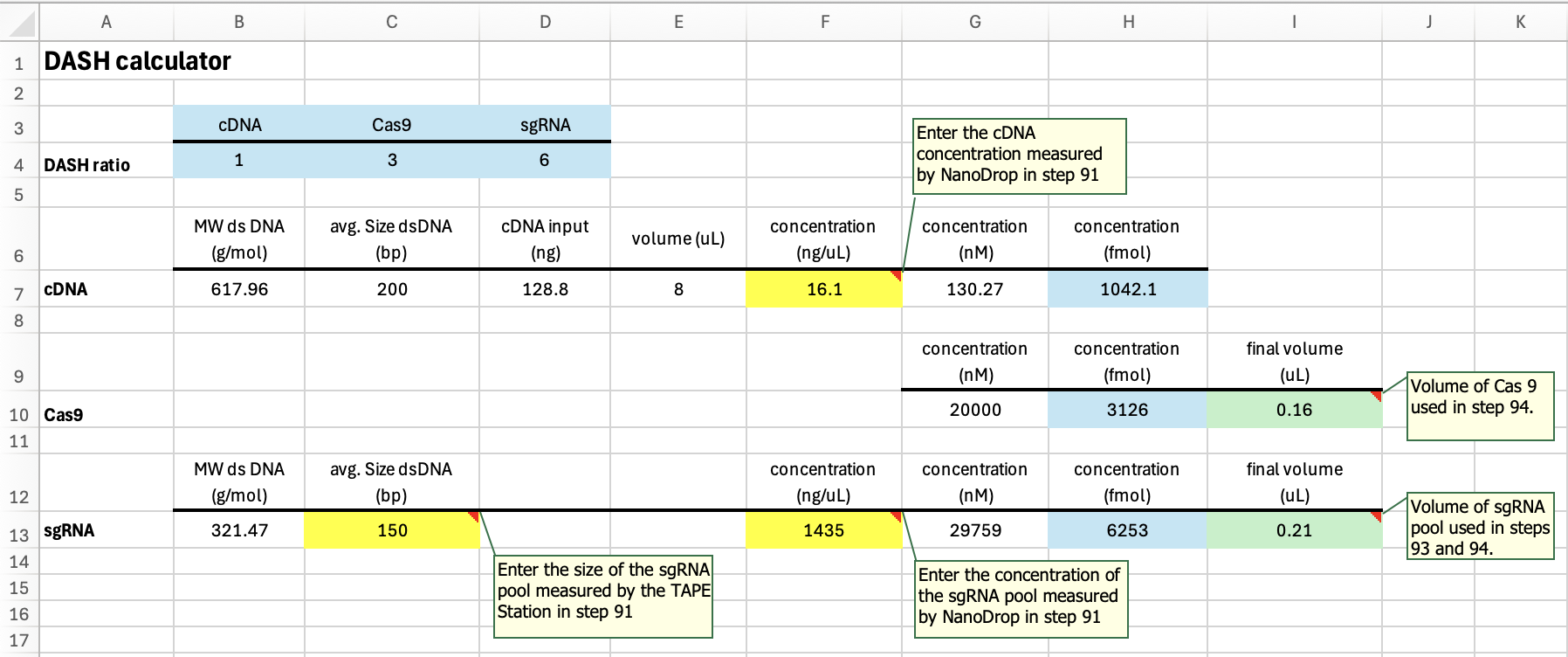

93| Add calculated sgRNA (ex. 0.21 μL) and nuclease-free water (amount per the table below) to a new PCR tube, incubate the tube at 94°C for 3 min, 4°C for 5 min.

94| Prepare the DASH reaction:

| **Reagent** | **Volume (****µL ) per reaction** | **Example reaction (µL)** |
| --- | --- | --- |
| sgRNA pool | per the calculation | **0.21** |
| Nuclease-free water | up to 10.5 | 8.28 |
| Cas9 buffer (3.1 NEB buffer) 10x | 1.85 | 1.85 |
| Cas9 | per the calculation | **0.16** |
| Mix total volume (after step 95) | 10.5 | 10.5 |
